# High-Quality Cell Capture Reveals Transcriptomic Changes After Single-Axon Injury of Mauthner Cells

**DOI:** 10.1101/2025.03.17.643681

**Authors:** Zheng Song, Lin Zhu, Yueru Shen, Huaitong Yao, Yuan Cai, Lingyu Shi, Xinghan Chen, Along Han, Ziang Zhao, Kun Qu, Bing Hu

## Abstract

In vivo single-cell capture methods based on micropipettes are essential for correlating morphological characteristics with transcriptomic profiling under physiological conditions. However, they often suffer from significant contamination of off-target cells. Consequently, we developed Hip-seq, which significantly reduces heavily contaminated cells, decreases contamination levels, improves cell reproducibility, and enhances data analysis accuracy compared to patch-seq. Using Hip-seq, we found that axon regeneration failure could be due to abnormal activation of translation and the circadian clock. Axon-regenerable central neurons reactivated axon development-related genes during regeneration. And one-to-one associated analysis helped identify pro-regenerative genes.

## Introduction

The rapid development of single-cell RNA sequencing (scRNA-seq) has significantly advanced the field of life sciences. Single-cell capture methods can be broadly categorized into three groups: suspension-based methods (such as FACS and 10x Genomics), micropipette (MP)-based methods (such as Patch-seq), and laser microdissection-based methods (such as LCM and fs-lm). Each method possesses unique advantages. Among these methods, MP-based single-cell capture allows direct capture of individual cells in vivo, preserving not only the physiological state of the cells but also integrating multiple aspects of cellular characteristics, including molecular, morphological, spatial, and physiological features[1, 2]. Furthermore, this method is not hindered by factors such as cell size and quantity, which limit suspension-based methods[3]. Due to these unique advantages, MP-based in vivo single-cell methods have gained significant traction among numerous researchers.

However, a common issue of scRNA-seq is contamination, which can lead to erroneous analysis results[4]. For example, contamination can lead to inaccurate identification of cell markers during single-cell subtype analysis and marker gene identification[5]. In differential expression analysis, contamination can lead to erroneous interpretations by erroneously including genes in the list of differentially expressed genes (DEGs). This is exemplified in an experiment using patch-seq to study axon regeneration, where the oligodendrocyte-specific expressed gene *Mbp* and red blood cell-specific expressed gene *Hbb-bs/Hba-a1* appeared in the list of DEGs[6]. These are typical marker genes that can be excluded based on experience, but more concerning is those nonspecifically expressed genes that can impact analysis results unnoticed. In suspension-based single-cell sequencing, several methods have been developed to mitigate contamination, such as Hydro-Seq[7]. Recently, a novel approach has been proposed for laser microdissection-based methods (fs-lm)[8], which effectively removes contaminants by washing after capturing intact cells. However, for MP-based methods, there is no method for removing contaminants, which has forced many researchers to halt their work[9, 10]. Certainly, numerous algorithms have been designed to mitigate contamination[11, 12]. However, it is important to note that regardless of the efficacy of remedial methods, they cannot be as effective as directly addressing the source.

To address the contamination issues in MP-based methods, we introduce the Hip-seq, a method for high-quality in vivo single-cell capture. Hip-seq enhances cell quality by removing off-target cells adhering to the outside of the MP and controlling the quality of the cDNA library before sequencing. Compared to patch-seq, Hip-seq effectively reduces the influence of contamination and improves the accuracy of data analysis. Using Hip-seq, we successfully captured single Mauthner cells (M-cells) from zebrafish larvae and obtained transcriptomic data. M-cells are a pair of neurons located in the hindbrain of each zebrafish[13]. In addition, their axons can regenerate after injury[14–16]. By analyzing our dataset, we found that failed axon regeneration might be caused by the activation of abnormal programs associated with translation and the circadian clock. And axon-regenerating central neurons reactivate axon development-related genes during axon regeneration. Moreover, we demonstrated that phenotype-associated analysis aids in identifying pro-regenerative genes. In summary, we developed a powerful method for single-cell capture in vivo and generated a distinctive dataset for identifying axon regeneration related genes in the CNS.

## Results

### 1. The workflow of Hip-seq for capturing high-quality single cells in vivo

Based on our experience, contamination in MP-based in vivo single-cell capture primarily stems from two sources: off-target cells adhering to the outside of the MP and off-target cell components introduced into the MP during cell aspiration. Therefore, Hip-seq primarily focuses on addressing these two primary sources of contamination to enhance cell quality. After the MP is tightly adhered to the target cell membrane, an appropriate negative pressure is applied using a syringe to aspirate the cell contents into the tip of the MP (the tip contains fluorescent dye indicating the position of the tips and a recombinant RNase inhibitor preventing RNA degradation). The tip was then immersed in 50 mM NaOH for up to ten times to completely remove adherent cells from the outer wall. Subsequently, full-length double-stranded cDNA libraries were generated according to the smart-seq2 protocol[17]. Following the exclusion of low-quality libraries, as determined by a bioanalyzer, 0.5 μl of each remaining sample is utilized for a second preamplification. This process generated sufficient templates for quantitative real-time PCR (qPCR). The relative expression levels of target and potential off-target cell-specific genes are then detected using qPCR. Samples with high expression of off-target cell-specific genes are discarded, and the remaining samples (the remainder of the first preamplification) are subjected to high-throughput sequencing (Fig 1A).

**Figure 1.**
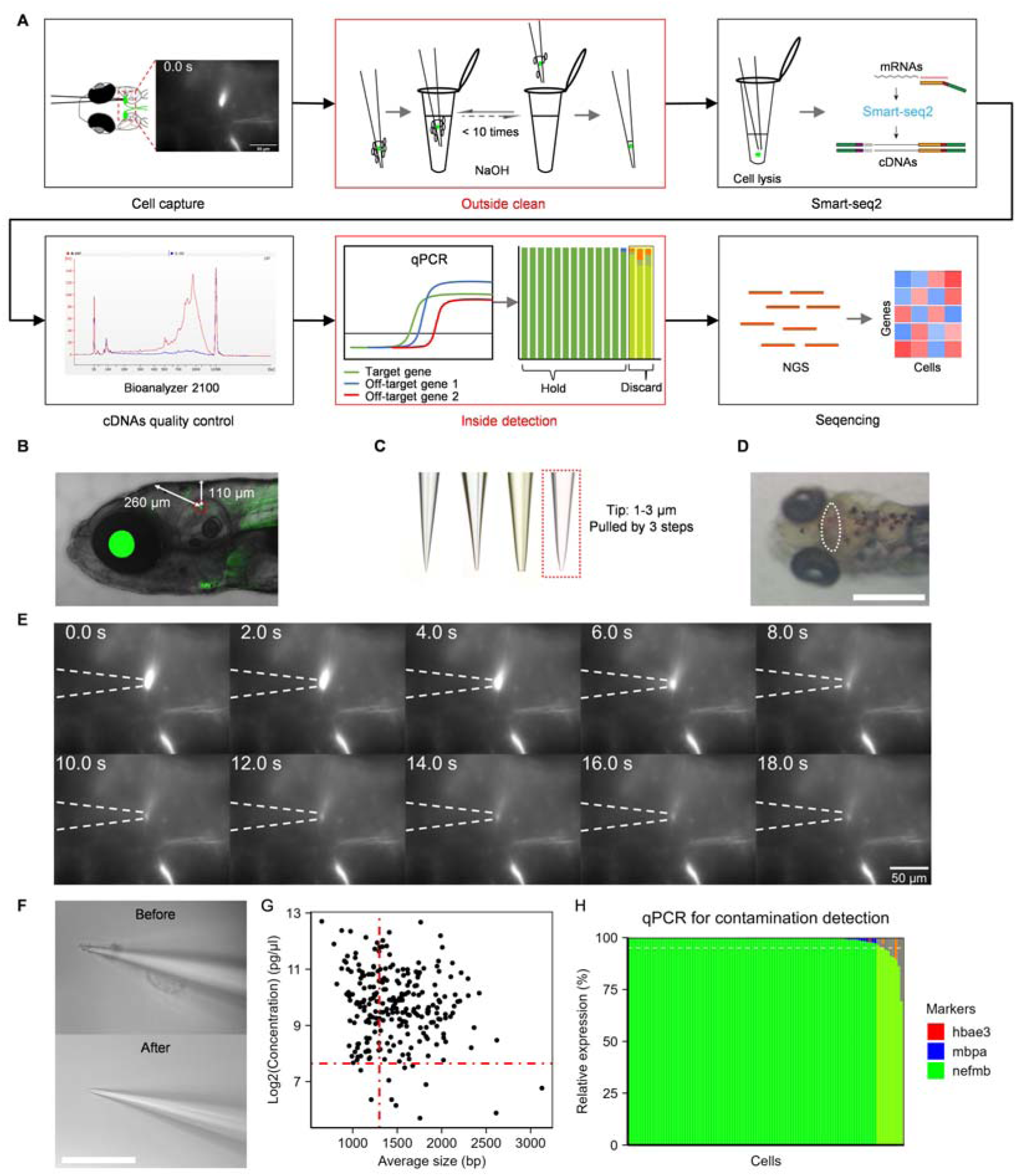
In vivo single-cell capture using Hip-seq. (A) Schematic diagram of Hip-seq. The red boxes represent the unique steps of Hip-seq. (B) Lateral illustration depicting the location of M-cells in the brain of zebrafish larvae. The center of the red dashed circle represents the position of the M-cell. The vertical and slanted bidirectional arrows indicate the distance between the M-cell and the brain skin (110 μm), as well as the distance which the micropipette passes through when capturing M-cells (260 μm), respectively. (C) Images of different micropipette tips. The red dashed box indicates the ideal tip for capturing M-cells, which were pulled using a three-step method and had a shorter taper tip diameter of 1-3 μm. The others represent micropipettes with small tip diameter, longer taper, and big tip diameter, respectively. (D) Dorsal illustration showing the tiny incision on the skin above the brain of zebrafish larvae. The white dashed ellipse represents the position of the incision. Scale bar, 500 μm. (E) Time-series schematic diagram depicting the capture of M-cells. Dashed lines indicate the position of the micropipette. Scale bar, 50 μm. (F) Representative images of tips before and after contamination removal with 50 mM NaOH solution. The top image represents the tips before contamination removal, while the bottom image represents the tips after contamination removal. Scale bar, 200 μm. (G) Scatter plot showing the average sizes (from 300-9,000 bp) and the concentrations of the captured M-cells detected by the bioanalyzer. The horizontal red dashed line indicates a concentration of 200 pg/μl, while the vertical red dashed line represents an average size of 1300 bp. (H) Stacked bar charts showing the relative expression ratios of genes specifically expressed in target and off-target cells. In this context, cells highlighted in bright yellow indicate a relative expression ratio of *nefmb* below 97%, resulting in their exclusion from further sequencing. n = 104 and 10 cells have a relative expression ratio of *nefmb* less than 97%.

### 2. Using Hip-seq to capture individual M-cells in vivo

To evaluate the performance of Hip-seq, we captured M-cells from zebrafish larvae. M-cells are located approximately 110 μm beneath the hindbrain of zebrafish larvae at 6 days post fertilization (dpf). However, due to the angled entry of the pipette into the brain, the actual distance traveled is approximately 260 μm (Fig 1B). To facilitate the aspiration of M-cell contents, we employed a three-step pulling method to fabricate the MP, resulting in a larger tip taper with a diameter of 1-3 μm (Fig 1C). To prevent the tip from being clogged when piercing the brain skin, a small incision was made at the entrance of the skin using ophthalmic scissors before the M-cells were captured (Fig 1D). When approaching the M-cells, appropriate positive pressure was applied to allow the internal solution to flow out slowly, preventing substances from entering the MP. The identification of M-cells was based on their distinctive morphological features and the transgenic signal (Video S1). In our experience, the aspiration of a single M-cell can be completed within 30 seconds (Fig 1E and Video S2). The successful capture of a M-cell depends on the disappearance of fluorescence signals under a fluorescence microscope. Then, we immersed the tips in 50 mM NaOH solutions up to 10 times to remove the off-target cells adhering to the outside of the MP (Fig 1F and Video S3). The entire capture process, from larval fixation to cell lysis, took approximately 10 minutes.

To investigate methods for removing adherent cells from the outer wall, we used DMEM, the nonionic detergent Triton X-100 (1% concentration), and one of the most common alkaline solutions, NaOH solution (5/50/500 mM). The results indicated that 50 mM NaOH was sufficient to effectively remove adherent cells from the outer wall (Fig S1A).

The integrity of the cDNA libraries was evaluated using a bioanalyzer. High-quality samples exhibited a main peak at approximately 2,000 bp and lacked small peaks under 500 bp (Fig S2A). Low-quality libraries were subsequently filtered out (Fig S2B). Occasionally, cDNA libraries with two high peaks were discarded, even though they displayed high concentrations[2, 18] (Fig S2C). In this study, we assessed the quality of the cDNA library based on the parameters of average size, concentration, and peak distribution. Specifically, we eliminated samples that had an average size less than 1,300 bp or a concentration less than 200 pg/μl or exhibited two distinct major peaks within the range of 300-9,000 bp (104/248 passes through, 41.9%) (Fig 1G).

Contamination may also be introduced into the MP during the extraction of target cells. Once off-target cells are inadvertently extracted into the MP, they become inseparable. To avoid the cost of sequencing these contaminated cells, we implemented a pre-sequencing quality control step to eliminate them. Although the genes specifically expressed in M-cells remain unknown, we detected an antibody, 3A10, which can label M-cells and a subset of other neurons in zebrafish[19]. Therefore, we selected one of its antigens, *nefmb*, as the gene specifically expressed in M-cells. For off-target cells, we selected *hbae3* and *mbpa* as markers because of their potential for encountering red blood cells and oligodendroglia while approaching and capturing M-cells. We converted the directly measured *cq* values from qPCR into relative expression ratios and discarded cells with an expression ratio of *nefmb* lower than 97% (94/104 passed through, 90.38%) (Fig 1H). The remaining cells were subjected to smart-seq2. It is worth noting that this threshold can be adjusted according to specific requirements. The relative expression ratios from the scRNA-seq and qPCR data exhibited a high degree of consistency, indicating the high reliability of the pre-sequencing quality control (Figs S3A-S3C).

### 3. Hip-seq significantly reduces the number and degree of contamination of cells

To evaluate the efficacy of Hip-seq in removing contaminants, we also employed patch-seq to capture M-cells (patch-seq n = 64; Hip-seq n = 94). First, we assessed the expression of marker genes for more cell types. The results showed that significantly fewer off-target cell-specific genes were detected in the cells captured by Hip-seq than in those captured by patch-seq (Fig 2A). We also analyzed two published patch-seq datasets[6, 20], and the findings demonstrated that severe off-target cell-specific gene expression was detected in both datasets, which indicated the presence of universal contamination in the patch-seq data (Fig S4A). Next, we utilized the relative expression levels of these target and off-target cell-specific genes to score these cells (see Methods for details). A higher score indicates a lesser degree of contamination by off-target cells. Compared to patch-seq, Hip-seq significantly reduced the number (only 6% of Hip-seq samples had scores below 97, while 77% of patch-seq samples did) and extent (average score for Hip-seq: 98.99, while patch-seq averaged only 71.29) of contaminated cells (Figs 2B and 2C). Furthermore, we observed a significant correlation among the number of genes, cDNA concentration, and score (Fig S4B). Further analysis revealed that the detection of more genes was associated with higher concentrations and higher contamination levels (Figs 2D and S4C-S4E). This result suggested that the greater number of genes detected via patch-seq may be influenced by off-target cell contamination. Furthermore, a GO enrichment analysis of unique genes identified via patch-seq revealed that these genes were related to off-target cells, such as vasculature, which supports our speculation (Fig 2E). Subsequently, we conducted differential expression analysis between the patch-seq and Hip-seq data. The results revealed a significant number of upregulated genes in the patch-seq data (1,328 upregulated genes, 486 downregulated genes, avg_log2FC > 0.4, p_val < 0.05), many of which were off-target cell-specific genes, such as glial cell-specific genes (*mbpa/mbpb/olig2*) and red blood cell-specific genes (*hbbe2/hbb1.3/hbae3/hbb1.1/hbae1.2/hbae1.3*) (Fig 2F).

**Figure 2.**
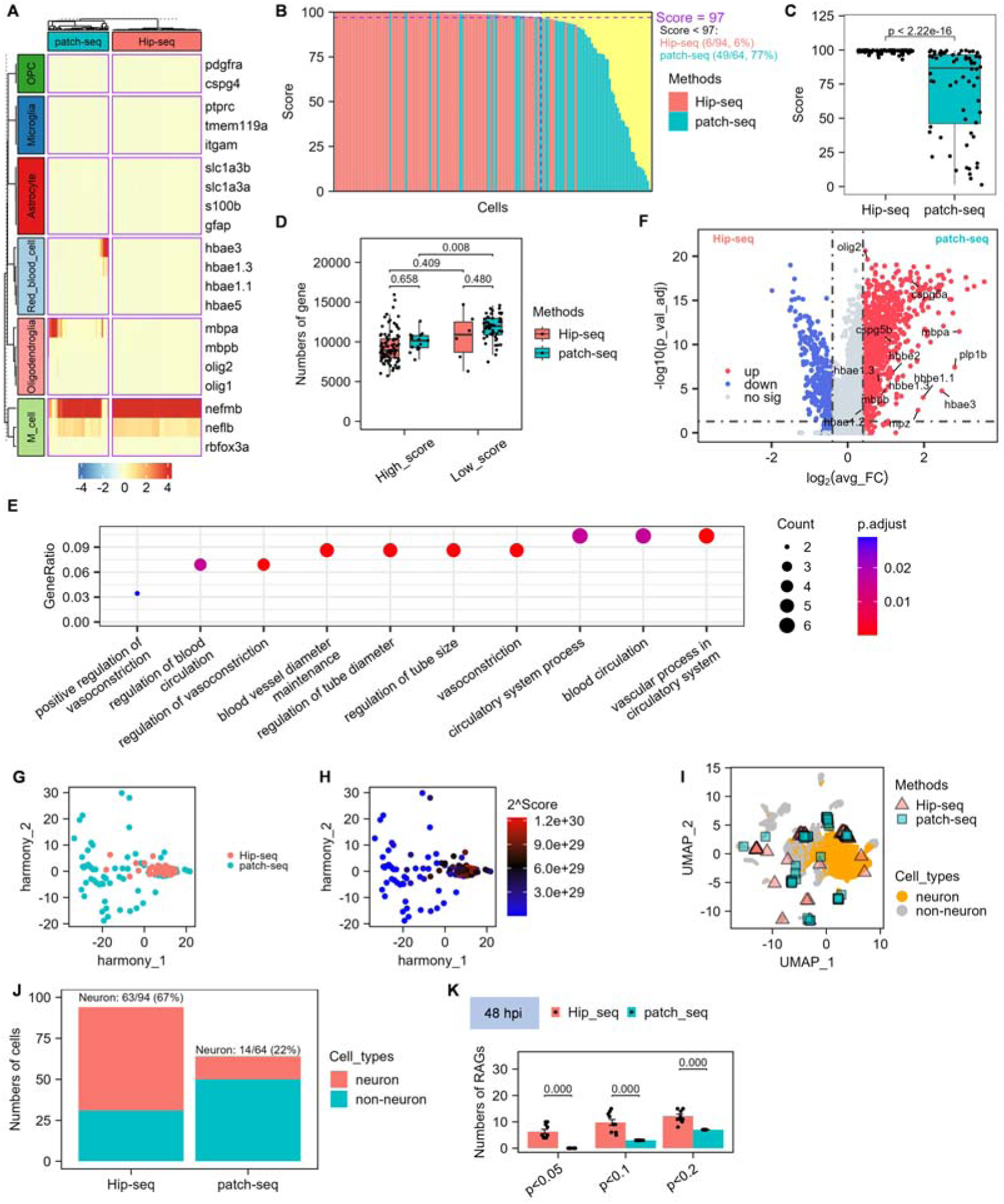
Hip-seq enhances cell quality. (A) The heatmap displays the expression profiles of marker genes of target cells and off-target cells captured by patch-seq and Hip-seq. (B) Bar plot showing the cell scoring. The scores decreased from left to right. The yellow region indicates cells with scores below 97. (C) Boxplots of cell scoring for patch-seq and Hip-seq. The box represents the interquartile range (IQR) between the first quartile (Q1) and the third quartile (Q3), with the median indicated by the line inside the box. Whiskers extend to the maximum and minimum values outside 1.5 times the IQR from Q1 and Q3, respectively. Statistical significance was determined using the two-sided Wilcoxon test, *p* < 2.2e-16. n = 64 for patch-seq, n = 96 for Hip-seq. (D) Boxplots of the number of genes detected in high-scoring and low-scoring cells. Cells with scores greater than 97 were classified as high-scoring, while the others were classified as low-scoring. The box represents the IQR between Q1 and Q3, with the median indicated by the line inside the box. Whiskers extend to the maximum and minimum values outside 1.5 times the IQR from Q1 and Q3, respectively. Statistical significance was assessed using two-way ANOVA followed by Tukey’s HSD test, with adjusted *p*-values indicated in the figure. Genes with TPM > 1 were included in the analysis. The average numbers were 9,394 (Hip-seq&High_score), 10,002 (patch-seq&High_score), 10,632 (Hip-seq&Low_score), and 11,804 (patch-seq&Low_score). (E) Top 10 GO terms of unique genes identified by patch-seq. (F) Volcano plot depicting differential expression analysis of cells captured by patch-seq and Hip-seq. Genes labeled in the plot represent off-target cell marker genes. The Wilcoxon test was employed for analysis. The horizontal dashed line represents *p* = 0.05, while the vertical dashed lines represent fold changes of 0.25 and -0.25. (G) PCA results of gene expression profiles for individual cells captured by patch-seq and Hip-seq. Batch correction between the patch-seq and Hip-seq data was performed using Harmony. (H) Relationship between cell scores and PCA distribution. The color represents the power of 99 of the scores. (I) Visualization of all cells using a UMAP plot. Triangles represent our cells, while dots represent single cells from the whole zebrafish dataset at 8 dpf sourced from Bushra Raj and colleagues[21]. We reannotated the annotated cell types into neurons or non-neurons. (J) Stacked bar chart illustrating the number of cells captured by patch-seq and Hip-seq that were predicted to be neurons and non-neurons. (K) Bar plot representing the number of RAGs among DEGs in 5 cells under axon-injured and uninjured conditions at 48 hpi, with *p*-value thresholds set at 0.05, 0.1, and 0.2. Hip-seq results are based on 10 random samplings of cells with successful axon regeneration. And patch-seq results were derived from 10 independent tests with identical samples. The list of RAGs is sourced from a review article[22]. Statistical significance was assessed using two-way ANOVA followed by Tukey’s HSD test, with adjusted *p*-values indicated in the graphs.

The results above, from a biological perspective, indicate that Hip-seq can effectively reduce the impact of off-target cell contamination.

### 4. Hip-seq improves cell reproducibility

Next, we evaluated the Hip-seq and patch-seq data from a data perspective. The principal component analysis (PCA) results indicated that the Hip-seq results exhibited tighter clustering than did the patch-seq results (mean distance of Hip-seq: 5.65, patch-seq: 22.72), suggesting an improvement in cell reproducibility (Figs 2G and S4F). To assess whether the dispersion of PCA correlated with off-target cell contamination, we examined the distribution of cell scores in PCA. As shown in Fig 2H, cells with lower scores exhibited a more dispersed distribution, indicating that the PCA distribution was indeed influenced by contamination. We also used whole-fish single-cell transcriptomic data as the reference data to predict the cell types of cells captured by patch-seq and Hip-seq (GEO accession: GSE158142, only using cells at 8 dpf and reannotated the clusters into two groups: neurons and non-neurons[21]). Hip-seq revealed a threefold greater proportion of neurons than patch-seq (Hip-seq: 67%, patch-seq: 22%), further suggesting that cells captured by Hip-seq are cleaner (Figs 2I, 2J and Table S1).

The above results demonstrate that Hip-seq effectively enhances the reproducibility of cells while reducing interference from off-target cells.

### 5. Hip-seq significantly improves the detection rate of regeneration-related genes

Contaminated data may lead to unknown influences on the results of the data analysis. Therefore, it is pertinent to investigate whether high-quality cells captured through Hip-seq can enhance the accuracy of the data analysis results. Both patch-seq and Hip-seq capture cells with axonal damage at different time points. To assess the advantage of the use of clean data, we performed differential expression analysis on each dataset and examined the detection rate of regeneration-associated genes (RAGs). The list of RAGs was compiled from a review article[22]. Due to the limited number of pre- and post-injury M-cells captured by patch-seq (5 cells in each group), we randomly selected 5 pre- and post-injury cells from the Hip-seq dataset to mitigate the impact of cell quantity on the credibility of differentially expressed genes (DEGs). This process was repeated 10 times. At 48 hpi, regardless of whether the *p* value was set to 0.05, 0.1, or 0.2, the percentage of RAGs detected by Hip-seq was significantly greater than that detected by patch-seq (Fig 2K). This result indicates that Hip-seq can mitigate contamination interference, thereby enhancing the accuracy of scientific investigations.

### 6. M-axons start to regenerate at approximately 2.5 hours after axon injury

Although the general process of axon regeneration in M-cells has been described [14], it is crucial to emphasize that regeneration does not commence immediately following axon injury. Instead, it first undergoes Wallerian degeneration, a process involving the gradual degradation of the axon distal to the cell body. The axon proximal to the cell body also experiences short-distance retraction prior to regeneration [23–25]. We hypothesize that the period between axon injury and the initiation of axon regeneration is a critical phase for cells to respond to axon injury and trigger axon regeneration programs. To determine the earliest time of axon regeneration, we utilized a light sheet microscope for continuous observation of the axon degeneration and regeneration process (Fig S5A). First, we performed unilateral axon ablation using a two-photon laser and recorded the time of axon injury in zebrafish larvae at 6 dpf. Subsequently, the larvae with unilateral axon ablation were immediately transferred to a light sheet microscope for further observation. Images were acquired every 3 minutes for 6 hours. Consequently, at approximately 2.5 hpi, the M-axons began to regenerate (Figs S5B, S5C and Video S4). Based on this time point, we categorized the axon regeneration process into two stages: the initial stage (before 2.5 hpi) and the regeneration stage (after 2.5 hpi).

### 7. Changes in the Transcriptional Profile During Mauthner Axon Regeneration

In this study, we utilized Hip-seq to capture individual Mauthner neurons at various stages of axon regeneration, including the initial stage with uninjured axons (N2) and injured axons (I2), as well as the regenerative stage with uninjured axons (N48), successfully regenerated axons (Re48), and unsuccessfully regenerated axons (NRe48) (Fig 3A). For the regeneration stage, we focused on 48 hpi, as this is a critical period during which distinct phenotypic changes occur during axon regeneration, such as varying regeneration lengths and failed regeneration. t-distributed stochastic neighbor embedding (tSNE) analysis revealed distinct clustering patterns, with Re48 cells forming a separate cluster, N2 and I2 cells located centrally, and N48 and NRe48 cells positioned in the upper right quadrant (Fig 3B). To investigate the changes in the transcriptional profile following axon injury, we conducted differential expression analysis. The results revealed that only 8 genes exhibited significant changes during the initial stage of axon regeneration (Fig 3C). We noticed that *tet3* and *hif1ab*, which are known as RAGs, were upregulated in this stage (Fig 3C). *tet3* facilitates the expression of various RAGs through DNA demethylation [26, 27], while *hif1ab*, a hypoxia-inducible factor, has been shown to promote axon regeneration under hypoxic conditions [28]. *tpra* is a member of the nuclear pore complex that controls the nuclear export of mRNAs and proteins [29]. *mgll* encodes a monoglyceride lipase, which participates in the hydrolysis of the endocannabinoid 2-arachidonoylglycerol (2-AG), thereby participating in signaling cascades [30, 31]. We also observed the downregulation of *dagla* (Avg_log2FC = -0.34, p_val = 0.09), a gene encoding a diacylglycerol lipase that participates in the synthesis of 2-AG [30, 32]. These results suggested that 2-AG could be an inhibitor of axon regeneration. Among the downregulated genes, *kcnj19b* encodes a potassium inwardly rectifying channel subfamily J member. A previous study showed that overexpression of the homologous protein *kir2a* significantly inhibited axon regeneration [33], suggesting that kcnj19b plays a similar inhibitory role in axon regeneration (Fig 3C). *snx4* has recently been identified as a synaptic protein [34], and its downregulation was consistent with the downregulation of synaptic assembly related genes previously reported in RGCs with axon regeneration [35, 36], suggesting a potential inhibitory role for *snx4* in axon regeneration (Fig 3C). *slc25a47a* is associated with mitochondria [37], while *eif3e* plays a crucial role in protein translation. However, the reasons for the downregulation of these two genes during the initial stage of axon regeneration remain unclear (Fig 3C). Overall, the differential expression analysis suggested that axon-regenerable central neurons initiate processes such as DNA demethylation, nucleocytoplasmic transport control, hypoxia sensing, and lipid metabolism/signaling pathways post axon injury, while processes related to neuronal activity and synapses are downregulated.

**Figure 3.**
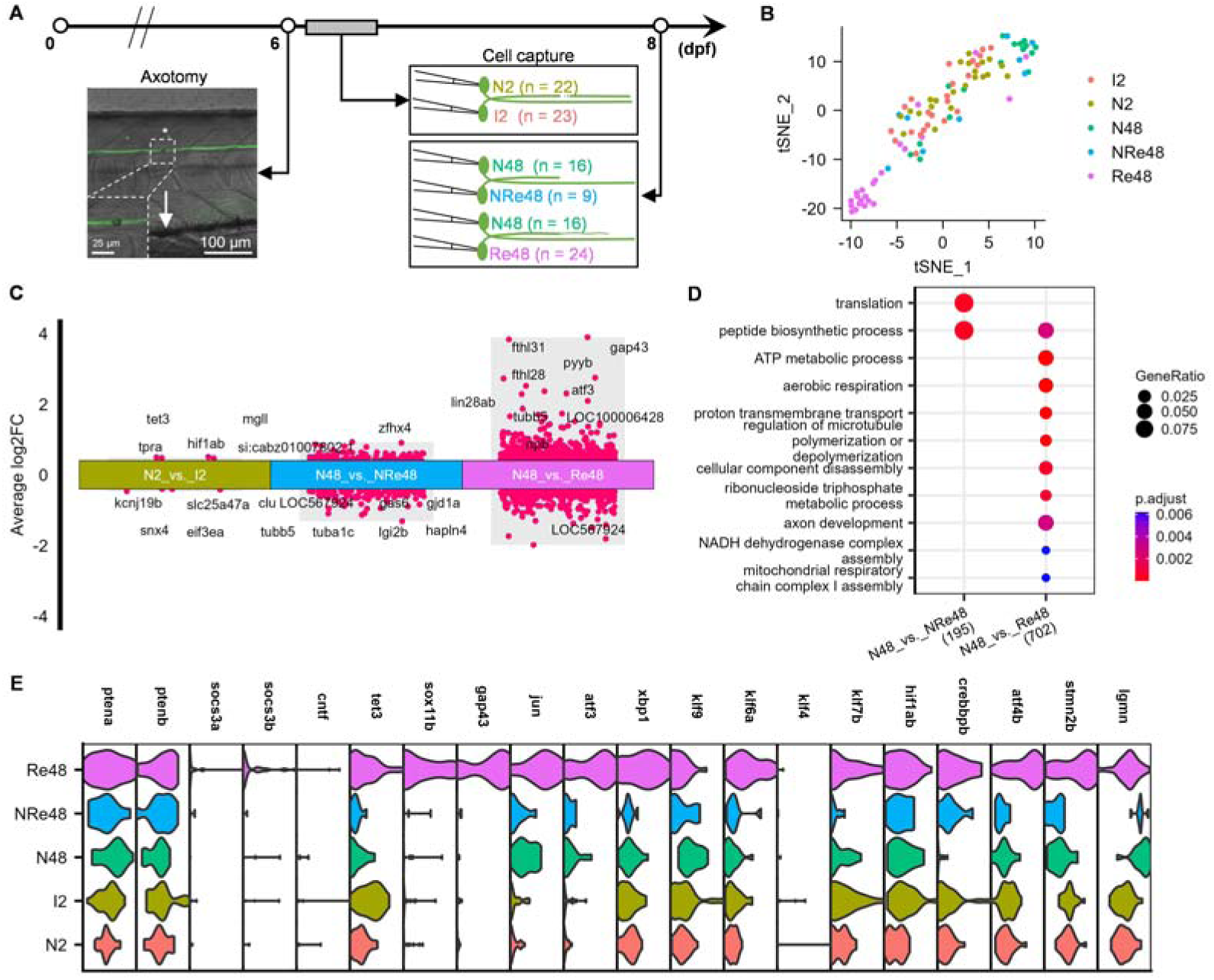
Gene expression profiles associated with M-axon regeneration. (A) Diagram illustrating the procedure for obtaining M-cells during the different stages of axon regeneration. N2 and I2 represent axon-uninjured and axon-injured M-cells at 50-150 min after axotomy, respectively. N48, Re48 and NRe48 represent axon-uninjured, axon regenerate successfully, and axon regenerate unsuccessfully at 48 hpi, respectively. n means sample number for each group. (B) The single-cell distribution after dimensionality reduction using tSNE. The different colors represent different groups. (C) Differential expression analysis of genes related to axon regeneration at the initial stage and regenerative stage. The labels are the top DEGs in each group. (D) Top GO terms for DEGs in each group. (E) Violin plots representing the expression patterns of classical RAGs in our dataset. The different colors represent different groups. The x-axis is the expression level.

Compared to those in the initial stage, a greater number of DEGs were differentially expressed between cells in the regenerative stage exhibited a greater number of DEGs whether with unsuccessfully or successfully regenerative axons (Fig 3C). To gain further insight into the biological processes associated with these DEGs, we conducted GO enrichment analysis. The results showed that DEGs of cells with unsuccessful regeneration of axons were enriched in translation and peptide biosynthetic processes, whereas DEGs of cells with successful regeneration of axons were enriched in processes such as peptide biosynthesis, energy metabolism, cytoskeleton organization, and axon development (Fig 3D).

Furthermore, we displayed the expression patterns of some classical RAGs in our dataset. Among them, the widely studied *ptena/b*, which has been a focal point in mammalian central axon regeneration research, did not exhibit significant changes during Mauthner axon regeneration. Additionally, genes such as *socs3a/b* and *cntf* exhibited minimal expression in Mauthner cells [35, 38, 39] (Fig 3E). This observation suggested that while these genes play pivotal roles in central axon regeneration, they do not undergo transcriptional regulation during axon regeneration in axon-regenerable central neurons. The expression of genes such as *sox11b*, *gap43*, *xbp1*, *atf3*, *jun*, *stmn2b*, and *atf4b* was exclusively upregulated in Re48, indicating that these genes are essential for successful axon regeneration (Fig 3E) [40–49]. Conversely, in NRe48, these genes remained inactive. The KLF transcription factor family also plays a crucial role in axon regeneration. The expression of proregenerative genes such as *klf6a* and *klf7b* was upregulated in Re48, while the expression of antiregenerative gene *klf9* was downregulated in Re48, consistent with their functions in axon regeneration (Fig 3E) [50]. However, another antiregenerative gene, *klf4*, was not expressed in Mauthner cells (Fig 3E). In addition to *klf9*, the gene *lgmn*, known to inhibit axon regeneration, was downregulated in Re48 (Fig 3E) [51]. Furthermore, *tet3* was significantly upregulated in I2 and Re48 but not in NRe48, suggesting a potential pivotal role in determining the fate of regenerating axons (Fig 3E). The expression of *hif1ab*, one of the earliest upregulated transcription factors during early regeneration, did not change in Re48, suggesting that it plays a predominant role in the initial response to axonal injury (Fig 3E). The expression of *crebbpb*, which promotes axon regeneration through p53 acetylation, was upregulated in both Re48 and NRe48 (Fig 3E) [52]. These findings suggest that in axon-regenerable central neurons, specific RAGs may regulate axon regeneration through pathways other than gene expression changes. The absence of *klf4* in Mauthner cells might be one of the crucial factors contributing to the regenerative capacity of Mauthner axons.

### 8. Activation of aberrant programs leading to axon regeneration failure

Previous studies have shown significant cellular heterogeneity in axon regeneration [35, 53, 54]. Using the single-axon injury model of Mauthner neurons eliminates cellular heterogeneity. Despite the strong regenerative capabilities of Mauthner axons, approximately 19% of these axons fail to regenerate (Figs 4A and 4B). Why do the same neurons exhibit contrasting outcomes in axon regeneration? Is it caused by a halted regeneration program or the activation of aberrant programs? The distinction between these two possible reasons is that the first can be reactivated with specific interventions, whereas the second requires additional steps to prevent aberrant activation.

**Figure 4.**
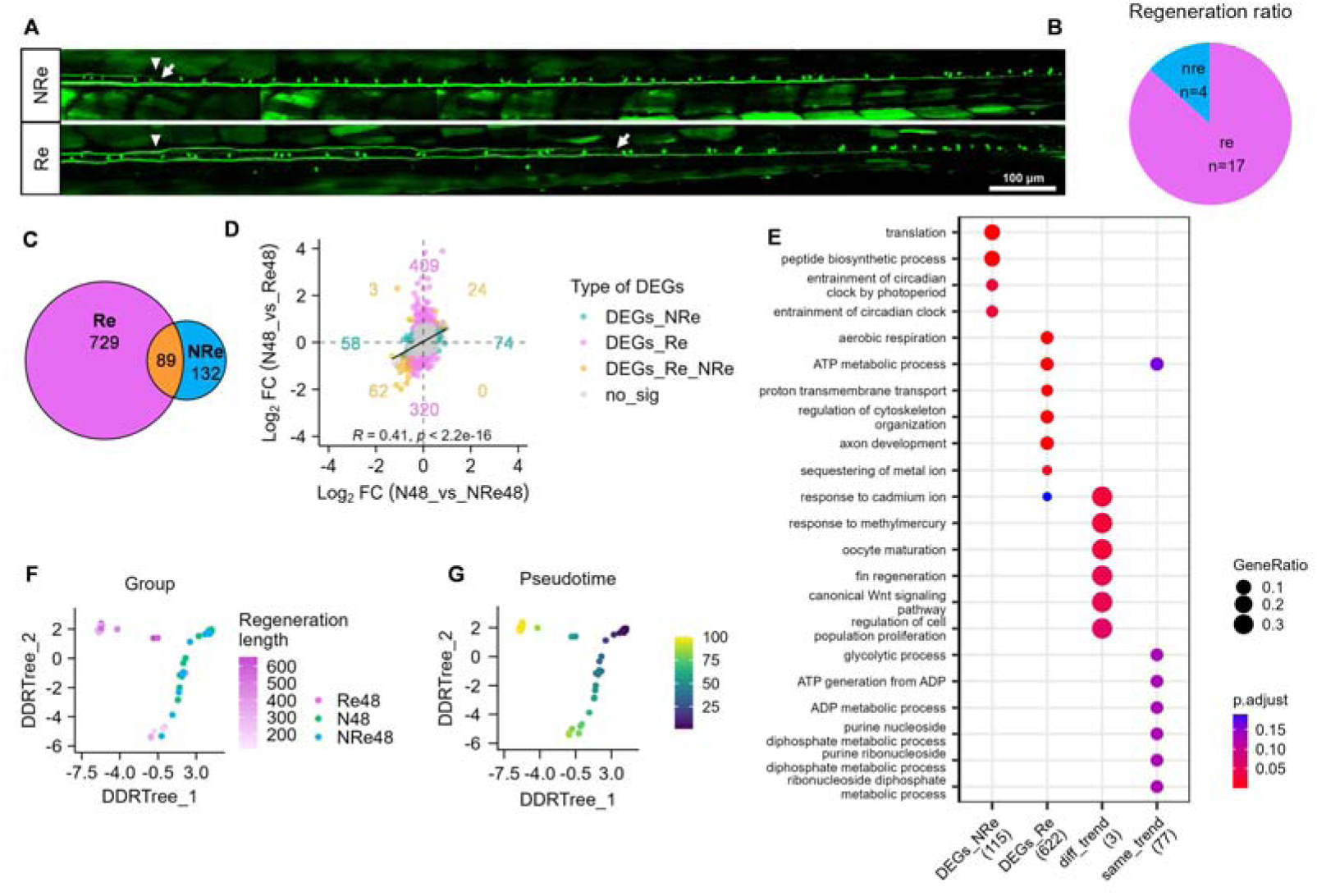
Activation of aberrant programs leads to axon regeneration failure. (A) Representative images of successful and unsuccessful axon regeneration. The white triangles denote sites of axon injury. The white arrows denote terminations of regenerative axons. Scale bar, 100 μm. (B) Regeneration ratio of M-axons. (C) Venn diagram showing the relationships between DEGs in Re48 and NRe48. (D) Scatter plots illustrating the Pearson’s correlation coefficients between genes in Re48 and NRe48 with p_val < 0.05. The colors represent the types of genes. The numbers represent the numbers of up- and downregulated genes of each type. (E) Top GO terms for DEGs in each group. DEGs_NRe indicates the unique DEGs in NRe48. DEGs_Re indicates the unique DEGs in Re48. diff_trend indicates DEGs with opposite change trends and shared by NRe48 and Re48, while same_trend indicates DEGs with the same change trends and shared by NRe48 and Re48. (F) The single-cell distribution after dimensionality reduction using DDRTree. Colors represent different groups, and transparency represents axon regeneration length. (G) Pseudotime trajectory analysis based on the dimensionality reduction results from DDRTree.

The GO analysis results above revealed that a unique term, translation, was enriched for DEGs in NRe48 (Fig 3D), suggesting that regeneration failure might be due to the activation of aberrant programs. To validate this hypothesis, we examined the relationships between DEGs in Re48 and NRe48. If this is true, there would be many unique DEGs in NRe48. As expected, approximately 60% of the DEGs in NRe48 were distinct from those in Re48 (Fig 4C). Consequently, we further examined the relationships among these DEGs. A scatterplot showed a moderate positive correlation in overall changes (Fig 4D). Although the majority of the shared DEGs exhibited similar patterns, three DEGs, namely, *tubb5*, *jun*, and *ddx5*, still displayed distinct trends (Fig 4D). *jun* was identified as the first transcription factor related to regeneration[44, 48]. We also examine the unique DEGs of Re48 and NRe48, and overlapped DEGs shared with same and different trends. The results showed that DEGs unique in NRe48 were associated with translation and circadian clock process (Fig 4E). And DEGs unique in Re48 were related to energy metabolism, cytoskeleton organization and axon development (Fig 4E). However, the overlapped DEGs shared with same trends were related to energy metabolism (Fig 4E). These results provide evidence that the failure of axon regeneration could be caused by abnormal activation of specific processes, potentially associated with abnormal translation and the circadian rhythm (details see in Table S3).

We also utilized Monocle to reconstruct the developmental trajectories between N48, NRe48, and Re48. If regeneration failure was a result of program interruption, NRe48 would be in an intermediate state. However, our findings showed that the developmental trajectory presented two branches (Figs 4F and 4G). Most of the N48 cells and a few NRe48 cells were positioned at the starting point of development, while the upper branch was mainly occupied by Re48 cells with stronger regenerative capacity (sRe48), and the lower branch comprised mostly NRe48 cells, along with a minority of N48 cells and some Re48 cells with weaker regenerative capacity (wRe48) (Figs 4F and 4G). This result further supports the hypothesis that regeneration failure is due to the activation of aberrant programs. Moreover, these findings indicate that cells with limited regenerative abilities share similarities with cells that fail to regenerate axons at the transcriptome level.

In summary, according to our results, we hypothesized that the failure of axon regeneration is attributed to the activation of abnormal programs.

### 9. Axonal development-related genes were reactivated during axon regeneration of axon-regenerable central neurons

In the preceding section, we observed that DEGs related to axon development were enriched in Re48 (Figs 3D and 4E). The regenerative capacity of axons gradually declines with age and disappears in adult mammals [55, 56]. Restoring the capacity for axon development in adult neurons is a method for stimulating axon regeneration [57–60].

From 6 to 8 dpf, Mauthner axons continue to grow along with ontogeny. To investigate whether axon regeneration requires the reactivation of the axon development process, N2 and N48 cells were considered two time points of axon development, while N48 and sRe48 cells were considered two states of axon regeneration. First, we examined the relationship between changes in gene expression during axon development and regeneration. Surprisingly, we found that genes significantly altered (p_val < 0.05) exhibited a moderate negative correlation overall (Fig 5A), which indicated that axon regeneration is an inverse process of axon development. This conflicts with the viewpoint that axon regeneration restarts genes related to axon development. Furthermore, we divided the DEGs into up- and downregulated DEGs and then performed separate GO enrichment analyses. The results suggested that genes associated with axon regeneration were upregulated during both axon development and regeneration. However, genes related to axon development were upregulated during axon regeneration but downregulated during axon development. (Fig 5B). Thus, the contradiction arose from axon development. Previous studies have suggested that Mauthner axons in zebrafish typically extend to the end of the spinal cord by 3 dpf [61]. Therefore, although axons continue to elongate slowly from 6 to 8 dpf along with body length [62], genes related to axon development, axon guidance, and synaptic transmission may no longer require high-level expression.

**Figure 5.**
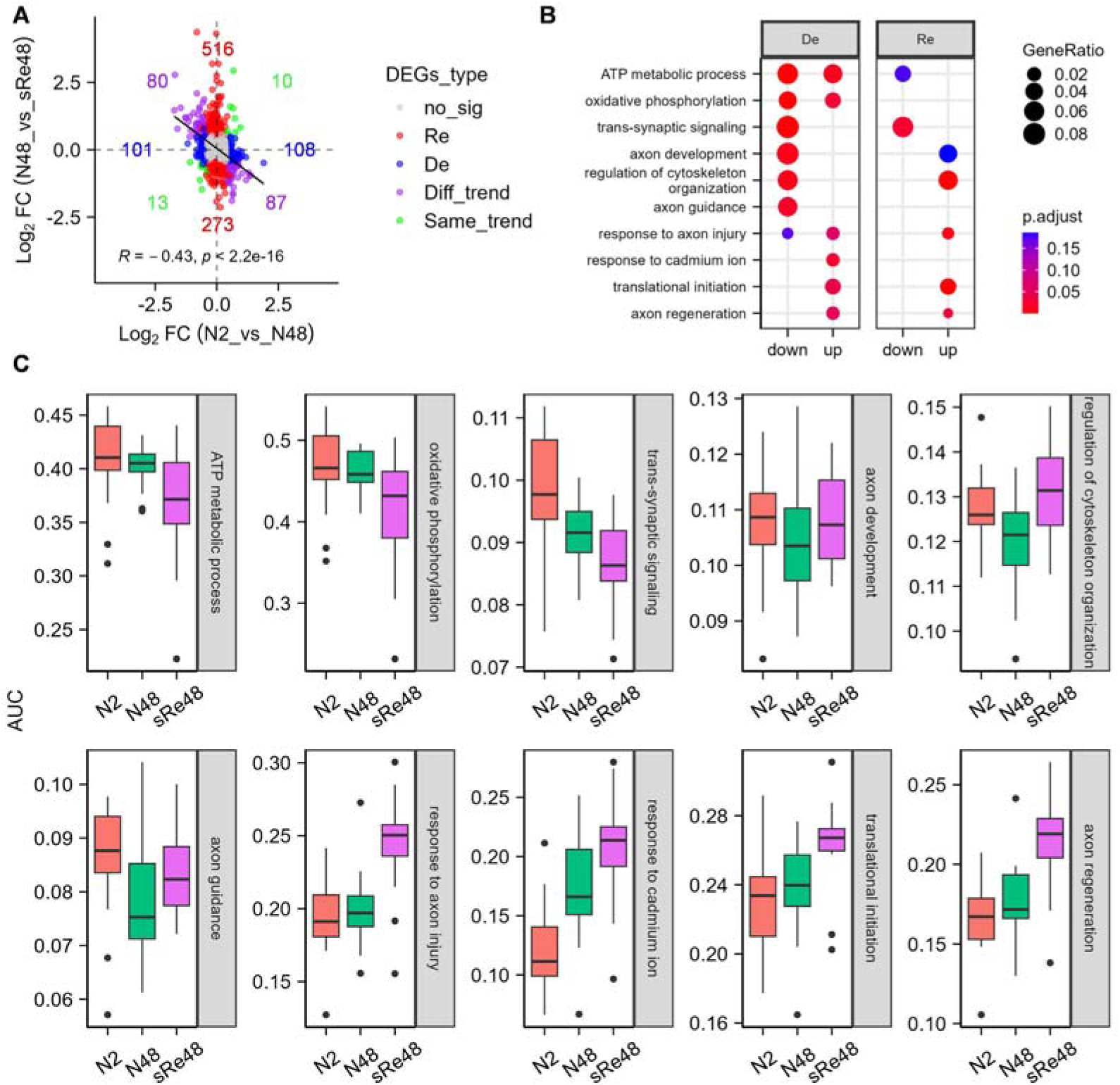
Axon development-related genes were reactivated during M-axon regeneration. (A) Scatter plots illustrating the Pearson’s correlation coefficients between genes related to axon development and regeneration with p_val < 0.05. The colors represent the types of genes. The numbers represent the numbers of up- and downregulated genes of each type. (B) Top GO terms for up- and downregulated DEGs in each group. De refers to axon development, and Re refers to axon regeneration. (C) Boxplots of the AUC determined by the AUCell of each group. The box represents the IQR between Q1 and Q3, with the median indicated by the line inside the box. Whiskers extend to the maximum and minimum values outside 1.5 times the IQR from Q1 and Q3, respectively. The dots represent outliers.

We also evaluated the biological activity of these biological processes using the AUCell algorithm. These findings corroborated previous results (Fig 5C). The above results suggest that axon-regenerable central neurons reinitiate axon development-related genes during axon regeneration.

### 10. Phenotype-associated analysis aids in identifying pro-regenerative genes

Identification of pro-regenerative genes is crucial for clinical treatment. One-to-one correspondence between each neuron and regeneration phenotype is a particular strength of the single-axon injury model of Mauthner neurons. To assess whether phenotype-associated analysis facilitates the identification of pro-regenerative genes, we evaluated the correlation between each gene and the regeneration length of 24 cells with successfully regenerated axons (Fig 6A). We detected a total of 45 negatively correlated genes and 84 positively correlated genes (cor > 0.4 or cor < -0.4, *p* < 0.05) (Fig 6B). The results from the GO analysis showed that the negatively correlated genes were associated with histone modification, Rho protein signaling transduction, and chemotaxis, while the positively correlated genes were related to protein catalysis, negative regulation of actin polymerization, and response to hypoxia (Fig 6C). These findings were consistent with previous research indicating that RhoA mediates the influence of extrinsic inhibitors of axon regeneration, such as CSPG, while HIF1A facilitates axon regeneration under hypoxia [28, 63]. Among these significantly correlated genes, we identified several known RAGs, such as *stmn2a* (also known as SCG10B, a classic marker of sensory axon regeneration) and *rhoab* (which inhibits axon regeneration) (Fig 6D) [63, 64]. Furthermore, we associated the correlation with DEGs to refine our screening scope for greater precision (Fig 6E). To validate the reliability of this strategy, we selected *tubb5* and *slc1a4*, the most significantly upregulated genes that are positively correlated with axon regeneration length, as candidates (Fig 6E). Using an in vivo single-cell electroporation technique, we overexpressed these two genes in individual Mauthner cells at 4 dpf and then damaged the axons at 6 dpf (Figs 6F-6H). By quantifying the length of regenerative axons at 48 hpi, we found that overexpression of *tubb5* and *slc1a4* significantly promoted axon regeneration (Figs 6I and 6J). The selection of these two genes is dependent on their expression patterns, indicating that phenotype-associated analysis can assist in the discovery of pro-regenerative genes.

**Figure 6.**
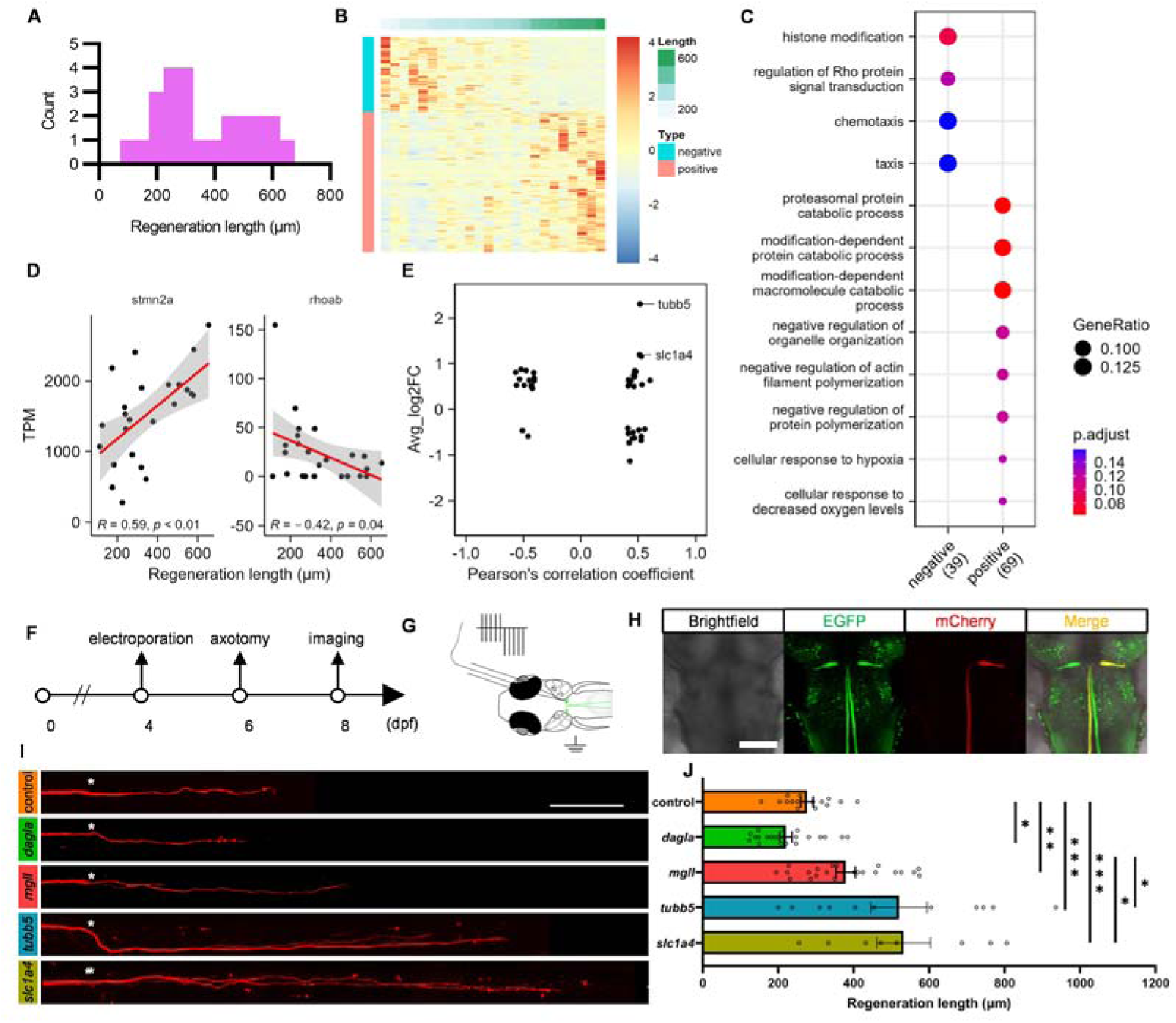
Phenotype-associated analysis aids in identifying pro-regenerative genes. (A) Distribution plot of axon regeneration lengths. n = 24. (B) Heatmap displaying genes negatively and positively associated with axon regeneration length. The top annotation indicates the axon regeneration length. (C) Top GO terms of negatively and positively associated genes. (D) Scatter plots illustrating the Pearson’s correlation coefficients between the expression levels of two RAGs and the axon regeneration lengths. The different colors represent different modules. Shaded areas represent the 95% confidence intervals around the means. (E) Scatter plots showing the relationships between fold change values (N48_vs_Re48) and Pearson’s correlation coefficients. (F) Flowchart illustrating the validation of candidate genes promoting the axon regeneration phenotype in the single-axon ablation model of M-cells. (G) Schematic representation of in vivo single-cell electroporation. (H) Representative images of zebrafish larvae with M-cells expressing the mCherry protein. The EGFP signal is from the endogenous signal of the transgenic zebrafish strain, indicating the positions of M-cells. The mCherry signal came from plasmids transfected into M-cells, indicating the successful expression of the target genes in M-cells. Scale bar, 100 μm. (I) Representative images of axon regeneration after overexpression of the candidate genes. White asterisks denote sites of axon injury. Scale bar, 100 μm. (J) Statistical chart for (I). Statistical significance was determined using two-tailed Student’s t tests. The average lengths were 276.375 μm (control), 220.455 μm (*dagla*), 378.682 μm (*mgll*), 520.091 μm (*tubb5*), and 532.75 μm (*slc1a4*). control, n = 16; *dagla*, n = 22; *mgll*, n = 22; *tubb5*, n = 11; *slc1a4*, n = 8. The data are presented as the mean ± SEM. * *p* < 0.05, ** *p* < 0.01, *** *p* < 0.001.

Our previous analysis suggested that the endogenous cannabinoid 2-AG may inhibit axon regeneration (Fig 3C), and we also tested *mgll* and *dagla*. The results demonstrated that overexpression of *mgll* significantly promoted axon regeneration, while overexpression of *dagla* inhibited axon regeneration (Figs 6I and 6J). We also noted that the pro-regenerative effect of *mgll* overexpression was significantly lower than that of *tubb5* and *slc1a4* (Figs 6I and 6J). Considering that *mgll* was significantly upregulated only in the initial stage, we hypothesized that *mgll* may play additional roles in signaling transmission.

In conclusion, these results demonstrate that phenotype-associated analysis facilitates the identification of pro-regenerative genes. Additionally, we displayed more genes that were predicted to take part in axon regeneration for further assessment (Fig S6).

## Discussion

In this study, we present a high-quality in vivo single-cell capture method based on MP, termed Hip-seq. This method enhances the quality of single-cell capture through two straightforward steps prior to sequencing. It can be integrated into existing methods to address the limitations of cell contamination, which allows researchers to confidently employ MP for in vivo single-cell capture. We further demonstrated that high-quality single cells can be captured without the need for a high-impedance sealing step, thereby reducing the complexity of the process. Compared to methods requiring cell suspension preparation, such as fluorescence-activated cell sorting (FACS) and droplet-based methods, Hip-seq reduces the cell capture time and minimizes transcriptional artifacts[65, 66] (from several hours to 30 s). In studies related to axon regeneration, the preparation of single-cell suspensions may initiate axon regeneration programs within a few hours following axon damage. This potentially introduces bias and hinders the detection of early-stage damage-specific changes. Hip-seq also preserves spatial, unique morphological and physiological features of cells, similar to patch-seq. Hip-seq is not influenced by cell size or quantity. While fs-lm is also unaffected by cell quantity[8], for larger cells, the required ablation point increases cubically with cell diameter. Additionally, Hip-seq does not require expensive two-photon lasers and can be implemented and utilized by a large number of researchers. In this study, we captured rare and large M-cells. In the future, Hip-seq could also be applied to in vivo studies of rare cells, such as adult neural stem cells and rare tumor cells, and studies related to neuronal transdifferentiation.

However, Hip-seq is labor intensive and has relatively low throughput. In the future, automation could reduce manual labor costs and enhance throughput.

We utilized Hip-seq for the first time to obtain the gene expression profiles of M-cells, which provides important reference data for Mauthner-related studies. Additionally, we explored the transcriptional alterations in M-cells following single axon damage at different stages. Unlike mammals, which require external interventions to promote axon regeneration, or *C. elegans*, which necessitate the use of transgenic strains to produce regeneration failure phenotypes, M-cells can exhibit both regeneration success and failure without any intervention[8, 14, 35]. Due to their rarity, M-cells effectively mitigate the impact of cell heterogeneity[35, 53]. Our results showed that the regeneration failure of M-axons could be caused by the activation of aberrant programs, such as translation and the circadian clock. However, this finding should be validated in the future. We also found that axon-regenerable central neurons reinitiated the axon development process during axon regeneration, which is consistent with previous studies[57–60]. Finally, our results showed the advantages of phenotype-associated analysis. In the future, comparisons of transcriptomic differences in axon regeneration between zebrafish and mammals could improve our understanding of central axon regeneration mechanisms.

In summary, we have developed the use of Hip-seq to address the issue of cell capture contamination. Furthermore, we have provided unique transcriptomic data on axon regeneration, thereby offering valuable insights into the intrinsic regulatory mechanisms of axon regeneration in the CNS.

## Conclusion

Hip-seq is an effective and translatable method for reducing off-target cells contamination in single-cell capturing in vivo. Using Hip-seq, we found axon regeneration failure could be due to abnormal activation of translation and the circadian clock, and identified several axon-regeneration associated genes. Anyway, our work provided a valuable dataset for axon regeneration in CNS, and Hip-seq will revive patch-seq related experiments that were previously abandoned due to contamination concerns.

## Materials and Methods

### Animals

All animal experiments conducted in this study were carried out in accordance with the guidelines and regulations set by the University of Science and Technology of China (USTC) Animal Resources Center and the University Animal Care and Use Committee (permit no. USTCACUC1103013). Zebrafish (*Danio rerio*) were maintained at a temperature of 28.5°C and subjected to a 14/10-hour light/dark cycle. Zebrafish embryos were collected in glass services and placed in incubators until further experiments. For larval anesthesia, ethyl 3-aminobenzoic methanesulfonate (MS-222, SigmaLAldrich, USA) at a concentration of 133-200 mg/L was used. Additionally, to prevent pigmentation, 0.003% N-phenylthiourea (PTU, SigmaLAldrich, USA) was added to the system water (0.2-0.3 g/L NaCl solution) between 24 and 48 hours post fertilization (hpf). Two strains of zebrafish were used in this study: AB and Tol-056. In this study, the zebrafish larvae used were of hybrid strains derived from the AB and Tol-056 strains. M-cells of the Tol-056 strain were labeled with enhanced green fluorescent protein (EGFP) obtained from RIKEN, Japan.

### Two-photon axotomy

To establish the single**-**axon ablation model, we used a two-photon laser to ablate individual M-axons. Specifically, after anesthesia, 6 dpf zebrafish larvae with labeled M-cells were immobilized using 1% low-melting agarose. Subsequently, the single M-axons were ablated using a Zeiss 980 two-photon laser (LSM980, Carl Zeiss, Germany) at a wavelength of 800 nm. The site of axon injury was above the cloacal pores in this study. Following axotomy, the larvae were released from low-melting agarose into the system water, which contained 0.003% PTU.

### Light-sheet microscope imaging

To observe the earliest time of axon regeneration, we employed live imaging to observe the process of axon regeneration with a light-sheet microscope (Luxendo MuVi SPIM, Bruker, Germany). Specifically, the axons of AB/Tol-056 larvae were ablated with a two-photon laser at 6 dpf, and the ablation time was recorded. Then, the larvae were immobilized in an FEP tube with 1% low-melting point agarose. Next, the FEP tube was installed in the sample chamber, and the sample chamber was filled with system water containing MS-222 and PTU. Images were captured at 3 min intervals continuously for 6 h using a 20×, 1.0 water-immersion objective lens. The captured images were converted to “x.ims” format files. Finally, videos of axon degeneration and regeneration were generated using Imaris software. The time at which the axon started to regenerate was analyzed from the videos.

### Capture the single M-cells in vivo

Micropipettes (TW100F-4, World precision instruments, USA) were pulled to capture single M-cells using a P-97 micropipette puller (Sutter, USA) employing a three-step method, resulting in a tip diameter of approximately 2-3 μm. This three-step method increased the tapering of the pipette tips, allowing for smoother capture of the cellular contents. Then, micropipettes were filled with approximately 0.3 μl of internal solution comprising 10 mM Alexa Fluor 594 (A10438, Thermo Fisher Scientific, USA) and 1 U/μl recombinant RNase inhibitor (2313A, Takara, Japan). Subsequently, the micropipette was mounted on a three-axis manipulator (MX7600, Siskiyou, Germany).

Zebrafish larvae of the AB/Tol-056 strain was immobilized with 2% low-melting agarose after anesthesia. Positioned dorsal side up on a culture dish covered with agarose, the larvae had their tails curled into a C-shape, aligning the two M-cells in the same horizontal plane, thereby facilitating the capture of both M-cells. The brain skin of the zebrafish larvae was manually peeled from the surface using ophthalmic scissors. It is crucial to avoid covering the larval brain with low-melting agarose during immobilization because agarose makes peeling difficult. Next, DMEM medium (NO. E600009, Sangon, Shanghai, China) supplemented with MS-222 (133-200 mg/L final concentration) was added to the culture dish to create an isotonic environment, and the larvae were maintained in an anesthetized state.

To prevent non-target cells and debris from entering as the tip approached the M-cells, we pushed a 1 ml medical syringe (approximately 0.05 ml) to generate positive pressure in the micropipette. This positive pressure allowed the internal solution to flow out slowly. The pipette was then adjusted to close the M-cell based on the signals from EGFP, Alexa Fluor 594, and brightfield under the water-immersion object (63x, 0.9 numerical aperture) of a fluorescence microscope. Subsequently, the pipette was adjusted to create a slight groove on the membrane of the M-cell. By pulling back half of a 1 ml medical syringe, negative pressure was generated to aspirate the cellular contents—typically sufficient to extract the cell contents.

Next, the micropipette containing the M-cell content was removed from the brain and dipped into 50 mM NaOH solution up to 10 times to eliminate off-target tissues and cells adhering to the outside wall of the micropipette. As a control, tips containing M-cells were not dipped in NaOH solution. Finally, the cell content from the micropipette was transferred to a PCR tube containing 4 μl of lysis buffer and placed on ice immediately. The lysis buffer comprised 0.1% Triton X-100 (No. A110694, Sangon, Shanghai, China), 2.5 mM each dNTPs (No. A610056, Sangon, Shanghai, China), 2.5 µM Oligo-dT30VN (5′-AAGCAGTGGTATCAACGCAGAGTACT30VN-3′, where ‘N’ is any base and ‘V’ is either A, C or G), and 1 U/μl recombinant RNase inhibitor (the primers used are listed in Table S2).

To prevent mRNA degradation, pipettes pulled on the same day were used. All water used in cell capture was RNase-free water, and the homemade tips for loading the internal solution into micropipettes were treated with 0.1% DEPC-water and autoclaved for 20 minutes. The liquid used during the cell capture process, such as the MS-222 solution added to DMEM, was filtered through a 0.22 μm filter. Maintaining a clean cell capture environment is crucial to avoid RNA degradation, as any minute solid particles or dust can lead to blockage of the micropipette tip, resulting in failure of cell capture.

### Test contamination removal reagents

DMEM, 1% Triton X-100, and 5, 50, and 500 mM NaOH solutions were utilized to evaluate the efficacy of micropipette outer wall adherent cells in removing contaminants. Following repeated immersion of the micropipette outer wall adherent cells in a PCR tube containing the test reagents 0, 1, 2, 5, and 10 times, the tips were observed under a microscope. While the DMEM, 1% Triton X-100, and 5 mM NaOH solutions were unable to completely eradicate the adherent cells, additional immersion in 50 mM NaOH 10 times was sufficient to completely remove them.

### Library construction and transcriptome sequencing

The RNAs extracted from individual M-cells were transformed into double-stranded, full-length cDNAs using the Smart-seq2 protocol, as previously described[17]. Subsequently, the cDNAs underwent 18 cycles of amplification using ISPCR primers.

The amplified cDNAs were then purified utilizing VAHTS DNA Clean Beads (N411-01, Vazyme, China). To evaluate the quality of the purified cDNA, 1 μl of the sample was analyzed using an Agilent 2100 Bioanalyzer (Agilent Technologies, USA) with a high-sensitivity DNA kit (5067-4626, Agilent, USA). Samples with low concentrations (typically less than 200 pg/μl) or short average sizes (usually less than 1300 bp within the 300-9000 bp range) were excluded. Prior to sequencing, cDNA samples passing the Agilent Bioanalyzer quality assessment were subjected to qPCR to determine the expression levels of cell-specific genes to identify and eliminate those with significant contamination. For this purpose, 1 μl of purified cDNA was subjected to an additional 18 cycles of amplification to serve as the template for qPCR (the primers used are listed in Table S2). In this study, the expression levels of *nefmb* (an M-cell marker), *mbpa* (an oligodendrocyte marker), and *hbae3* (a red blood cell marker) were detected. The cq values of each gene were then transformed into a relative expression ratio according to the following equation:

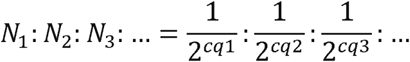

Here, *N_1_*, *N_2_*, and *N_3_* represent relative expression ratios, and *cq1*, *cq2*, and *cq3* represent the *cq* values obtained from qPCR analysis. Samples with a *nefmb* expression ratio less than 97% were excluded, and the remaining samples were subjected to RNA sequencing. For library preparation, the TruePrepTM DNA Library Prep Kit V2 for Illumina (TD503, Vazyme, China) was used, with 1 ng of purified cDNA used as input. Briefly, cDNA was fragmented and tagged using Tn5 transposase. The resulting tagged fragments were subjected to 15 cycles of PCR amplification. Fragments of approximately 200 bp were selectively isolated using VAHTS DNA Clean Beads. Subsequently, the prepared sequencing libraries were subjected to paired-end sequencing on an Illumina NovaSeq 6000 platform. To ensure data quality, low-quality reads were filtered out using fastp. The remaining cleaned reads were aligned to the zebrafish ribosomal RNA (rRNA) library with Bowtie2 to eliminate potential rRNA reads. Unmapped reads were then aligned to the zebrafish reference genome (GRCz11) using HISAT2. RSEM was used for gene expression quantification, and expression levels were normalized to transcripts per million (TPM) values.

### Cell scoring

Cell scoring is achieved by utilizing the TPM values of M-cells and non-target cell-specific expressed genes. The calculation is based on the following formula: Score = 100% * (Sum of TPM values of M-cell expressed genes) / (Sum of TPM values of M-cell and non-M-cell expressed genes). The genes utilized for this calculation are illustrated in Fig 2A.

#### Mapping to reference

Our data were mapped to the reference data (8 dpf data of GSE158142) using the “FindTransferAnchors” and “MapQuery” functions. The cell types were into neurons and non-neurons, based on existing knowledge (see Table S1).

#### Pseudotime trajectory analysis

The Monocle package was used for pseudotime trajectory analysis. DEGs were utilized as sorting genes. The spatial dimensionality was reduced to a two-dimensional space using “DDRTree”, and the “plot_cell_trajectory” function was applied to visualize the trajectory in the reduced-dimensional space.

#### Differential expression analysis

The Seurat package (v4.3.0) was used to construct a Seurat object using the count matrix in R. The resulting object underwent processing with the default parameters for normalization and ScaleData. Subsequently, DEGs were identified using the “FindMarkers” function with the Wilcoxon test. In this study, genes with a p_val less than 0.05 and an absolute avg_log2FC greater than 0.4 were considered DEGs, with the min.pct threshold set to 0.25.

#### Gene enrichment analysis

To explore the biological functions and pathways associated with the gene sets of interest, we conducted GO enrichment analyses for genes of interest with the functions “enrichGO” and “compareCluster” in the R package clusterProfiler. We set the method for adjusting the *p* value to “BH”. The function “simplify” of clusterProfiler was used to remove redundant terms.

#### AUC Scoring

To evaluate the biological activity of interesting biological processes, we extracted specific gene sets using the “get_GO_data” function of clusterProfiler. All genes of each gene set were used to calculate the area under the curve (AUC) scores using the R package “AUCell”.

#### Construction of overexpression plasmids

The overexpression system used in this study consisted of CMV-Gal4 and UAS-mCherry plasmids. The CDSs were cloned from a cDNA library of zebrafish and then inserted between UAS and mCherry sequences.

#### Single cell electroporation

To assess the impacts of candidate genes on axon regeneration, overexpression plasmids for these genes were constructed and delivered into individual M-cells through in vivo single-cell electroporation (the primers used are listed in Table S2). Initially, 4 dpf zebrafish larvae were anesthetized with MS-222 and immobilized using 1% low-melting agarose in a custom-made electroporation chamber. Subsequently, a pipette containing plasmids and Alexa Fluor 594 was gently positioned against the membrane of M-cells. For plasmid transfection, a series of sinusoidal wave pulses were applied via a single-cell electroporation system (Axon Axoporator 800A, Molecular Devices, LLC, USA), which created minute pores in the cell membrane, as previously described[33, 67]. Specifically, a 0.5-second alternating current pulse with a voltage range of -8 V to 8 V was employed. Next, a 0.5-second negative pulse with a voltage range of 0 V to -16 V was applied. Both the alternating current pulse and the negative pulse had a frequency of 200 Hz and a duty cycle of 10%. The concentration of each plasmid used in this experiment was approximately 100 nM. At 2 days post electroporation, larvae expressing the mCherry protein were subjected to axotomy to investigate the effects on axon regeneration.

#### In vivo imaging

To quantify axon regeneration lengths, in vivo imaging of regenerated axons was conducted using a laser scanning confocal microscope (FV1000, Olympus, Japan). Imaging was performed at 48 hpi. Imaging was performed with 40× and 0.85× water-immersion objective lenses. The scanning interval along the z-axis was set to 3 μm. We performed “intensity projection over the z-axis” and “save display” for each image in the “olympus fluoview” software (Ver 4.2b). Subsequently, the images were stitched together in lightened blend mode using Photoshop software (Adobe Photoshop 2020). The lengths of regenerative axons, defined as the straight stretch from the injury site to the terminal, were measured using FIJI software.

#### Statistical analysis

Axonal regeneration lengths between two groups were analyzed using unpaired two-tailed Student’s *t* tests with GraphPad Prism 9.0 software. The axon regeneration length data were tested for a normal distribution, and all of the data were normally distributed. The other statistical analysis methods used are listed in the figure legends.

## Acknowledgments

We thank Ji Tang for the help in imaging using light-sheet microscope. We also thank Xuyuan Gao for the help in the usage of bioanalyzer. This work was funded by the “Research Funds of the Center for Advanced Interdisciplinary Science and Biomedicine of IHM” (QYZD20220002), the National Natural Science Foundation of China, No. 82071357, and the Ministry of Science and Technology of China, No. 2019YFA0405600 (to B.H.).

## Supporting Information

**S1 Fig.**
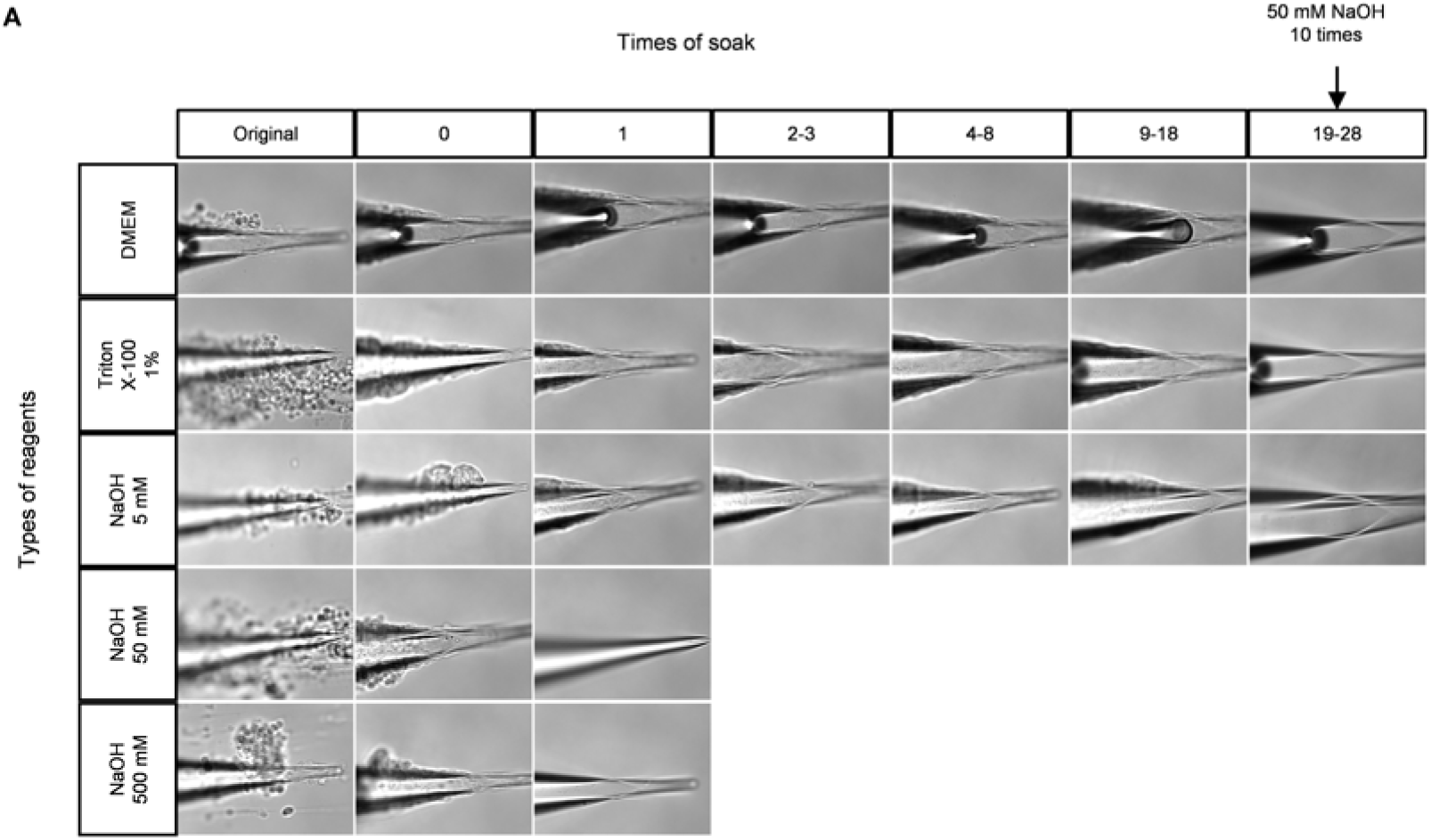
Comparison of different contamination removal reagents. (A) Results of different contamination remove reagents, including DMEM, 1% Triton X-100, 5 mM NaOH, 50 mM NaOH and 500 mM NaOH. Top annotation means the times of soak and left annotation means the type of reagents. When testing DMEM, 1% triton X-100 and 5 mM NaOH solution, even 18 times of soak cannot remove the off-target cells adhered to the outside of MP. However, following extra 10 times of soak with 50 mM NaOH, the remaining off-target cells can be removed cleanly. While testing 50 mM and 500 mM NaOH solution, sometimes first soak can remove them. But in our experience, up to 10 time can remove any cells of outside.

**S2 Fig.**
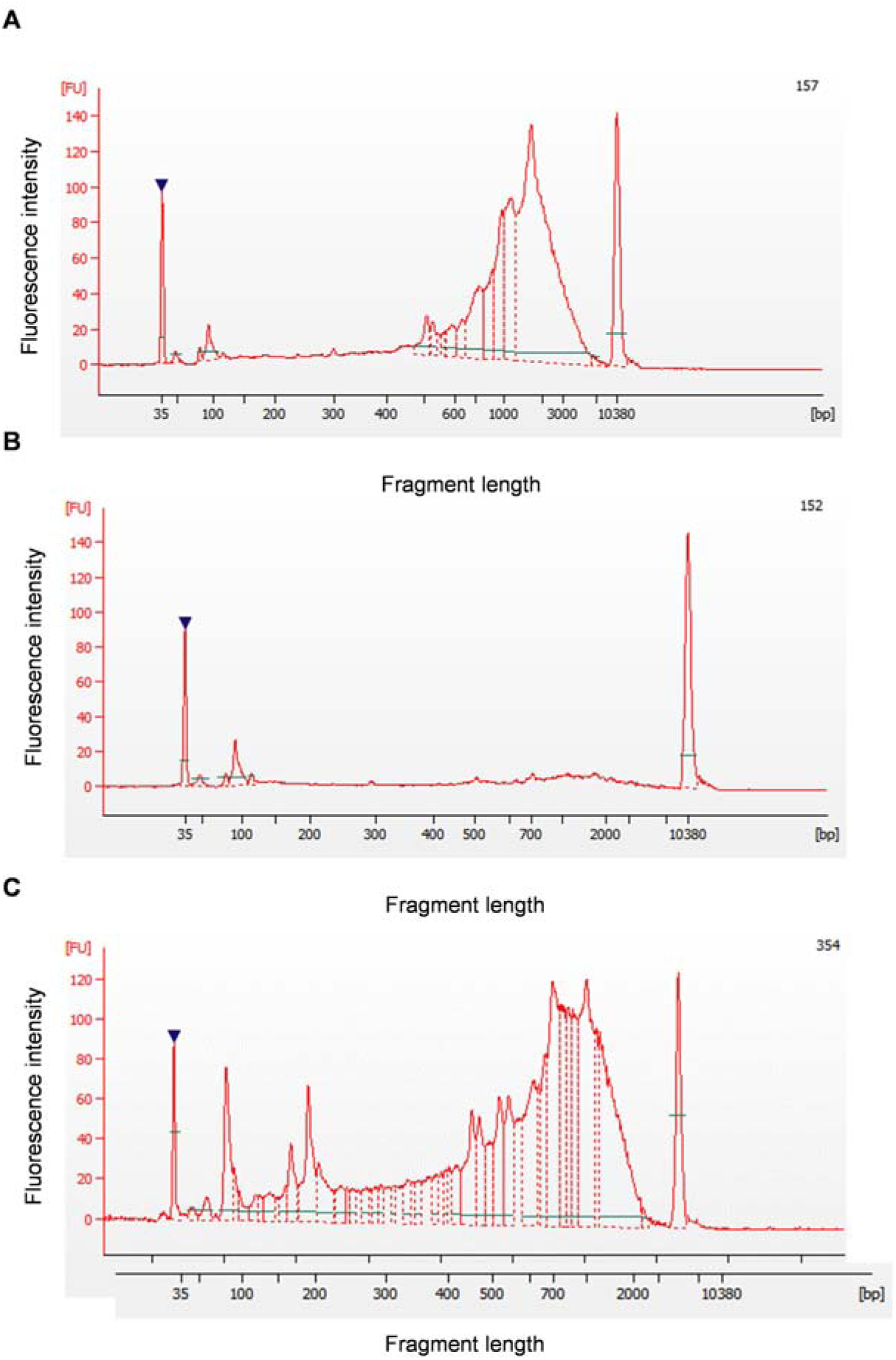
Quality control of amplified cDNA libraries. (A) An example of libraries of a good quality library. (B) An example of a severe degeneration library. (C) An example of a library with a high concentration and large average size but with two high peaks.

**S3 Fig.**
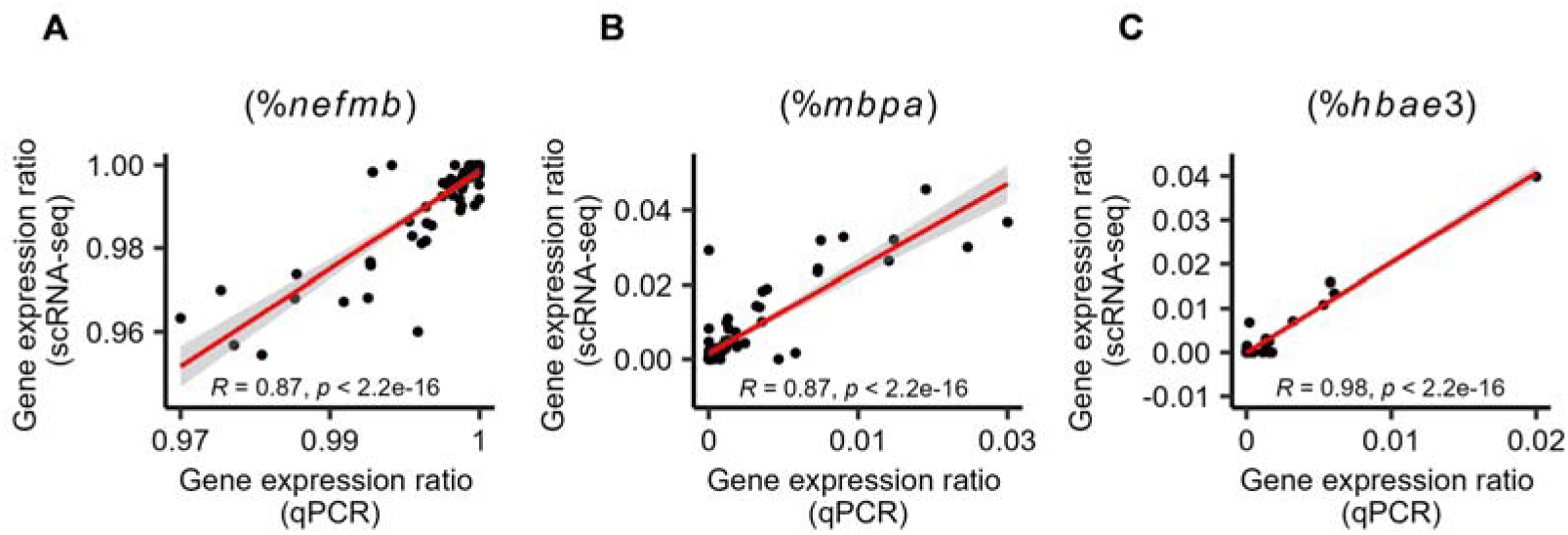
The accuracy of qPCR as the quality control method before sequencing. (A-C) Scatter plots depicting the Pearson’s correlation coefficient between gene expression ratios obtained through qPCR and scRNA-seq. (A) Correlations for the *nefmb* gene, (B) for *hbae3*, and (C) for *mpa*. Shaded areas represent the 95% confidence intervals around the means.

**S4 Fig.**
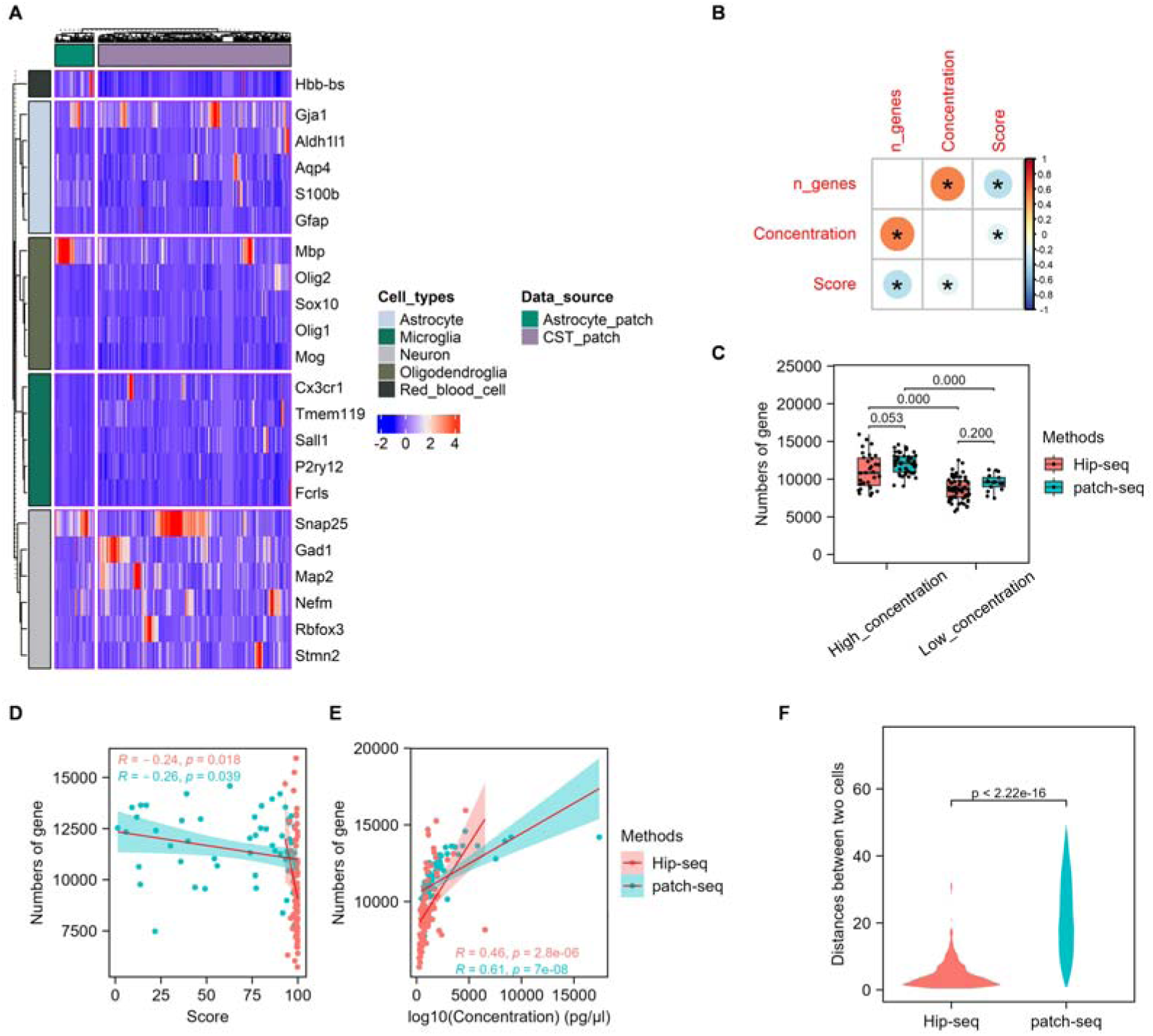
Hip-seq enhances cell quality. (A) Heatmap illustrating the expression of neuron and non-neuron marker genes. The data were sourced from single cells captured by patch-seq of two published articles^6,23^. (B) Heatmap depicting the correlations between the score, concentration, and number of genes. Cells are sourced from patch-seq and Hip-seq. The Pearson correlation coefficient was utilized. * indicates *p* < 0.05. (C) Boxplot illustrating the number of genes detected at different cDNA library concentrations. Libraries with concentrations above 1000 pg/µl were defined as having a high concentration, while those with concentrations below 1000 pg/ul were defined as having a low concentration. The box represents the IQR between Q1 and Q3, with the median indicated by the line inside the box. Whiskers extend to the maximum and minimum values outside 1.5 times the IQR from Q1 and Q3, respectively. Statistical significance was assessed using two-way ANOVA, with adjusted *p* values indicated in the graph. Genes with TPM > 1 were included in the analysis. The average numbers were 11,091 (Hip-seq&High_concentration), 12,047 (patch-seq&High_concentration), 8,677 (Hip-seq&Low_concentration), and 9,545 (patch-seq&Low_concentration). (D) and (E) Scatterplots illustrating the relationship between the number of genes and cell scores and between the number of genes and cDNA concentration, respectively. The red line segments represent the linear regression results, with the shaded areas representing the 90% confidence intervals. The numerical values in the graph indicate the Pearson correlation coefficient and *p* value. (F) Violin plots depicting distance statistics from harmony plots of cells captured by patch-seq and Hip-seq. Statistical significance was assessed using the two-sided Wilcoxon test.

**S5 Fig.**
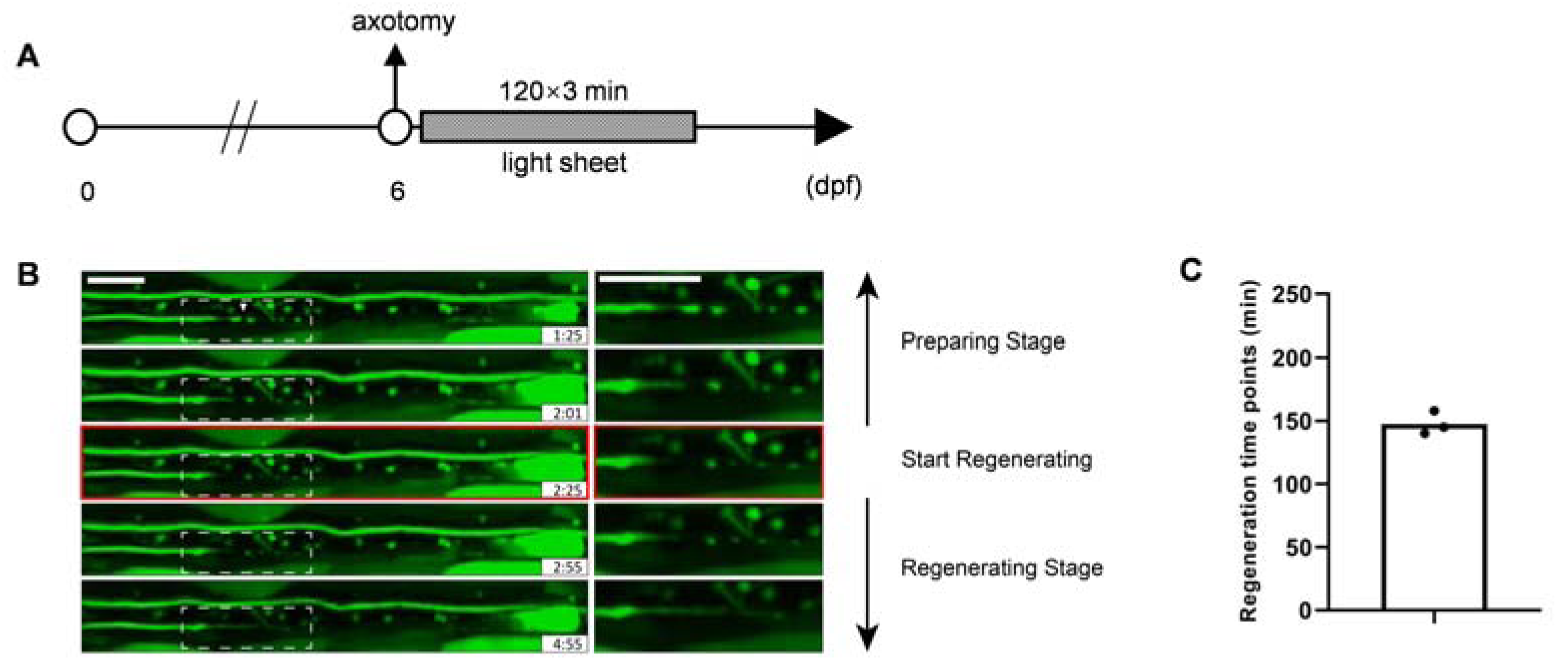
Observation of earliest time for M-axons to regenerate. (A) Experimental workflow diagram for observing the earliest time for M-axons to regenerate. Unilateral ablation of the M-axons was performed using a two-photon laser at 6 dpf. Subsequently, immediate observation was conducted using a light-sheet microscope, capturing images at 3-minute intervals for 6 hours. (B) Schematic diagram of a single axon undergoing two-photon ablation. The asterisk indicates the site of axon injury, and the arrow points to the location of the cloacal pores. The image on the right is an enlarged view of the dashed box on the left. Scale bar on the left, 100 μm. Scale bar on the right, 25 μm. (C) A time-lapse image series of axon regeneration, with the time in the lower right corner representing the post-injury time. The white triangle in the top panel indicates the site of axon injury. The images on the right are enlarged views of the white dashed box on the left. The phase before the onset of axon regeneration is defined as the preparatory stage, and the phase after the onset of regeneration is defined as the regenerative stage. Scale bar, 50 μm. (D) Bar plot depicting the onset time of axon regeneration.

**S6 Fig.**
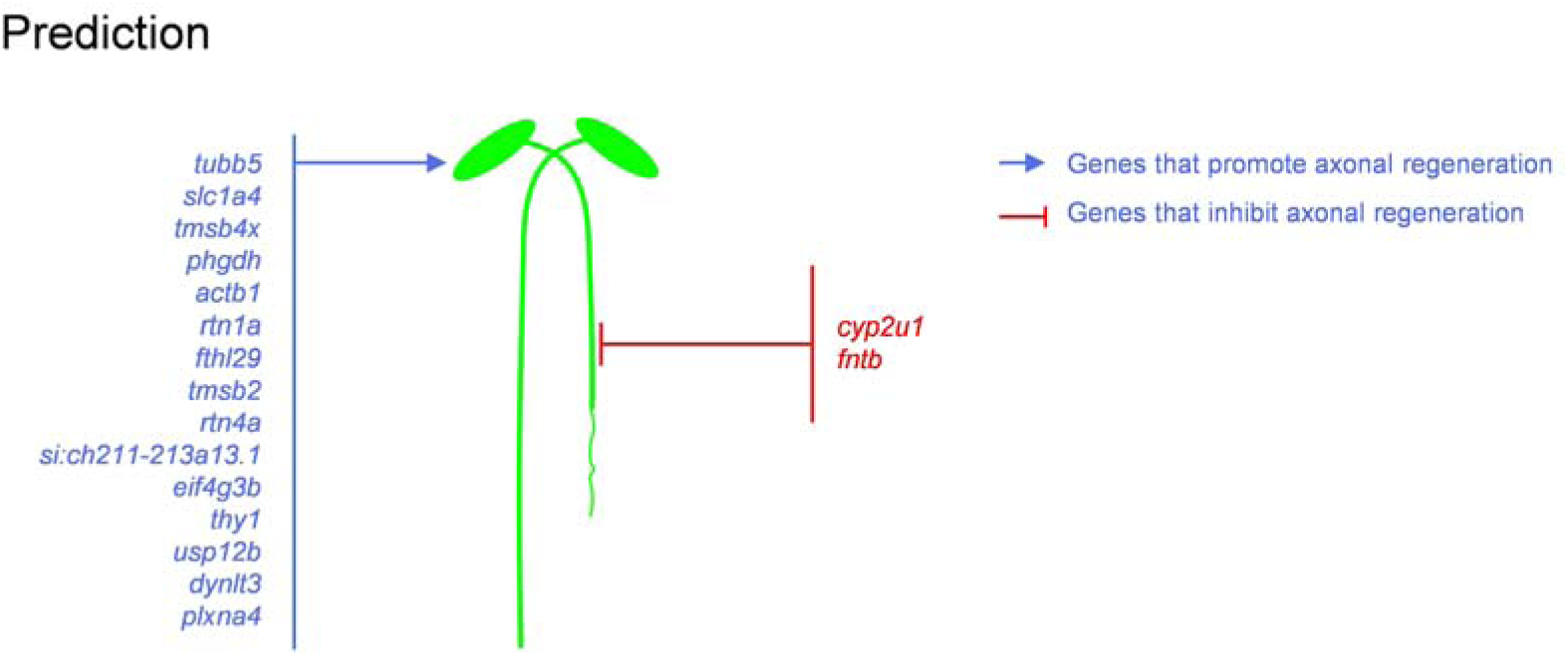
Prediction of genes that promote or inhibit axon regeneration. Blue represents genes that were predicted to promote axon regeneration, which were up-regulated in M-cells with successfully regenerated axons and were positively correlated with the regeneration length (cor > 0.4). Conversely, red represents genes that were predicted to inhibit axon regeneration, which were down-regulated in M-cells with successfully regenerated axons and were negatively correlated with the regeneration length (cor < -0.4).

## Data and code availability

The raw data and processed data obtained through patch-seq and Hip-seq in this study can be accessed through Gene Expression Omnibus (GEO) (accession number: GSE260744). Other data used in this study can be obtained through the following links: Astrocyte patch (https://doi.org/10.5281/zenodo.7704838); CST_patch (GSE205769); whole-fish single-cell transcriptomic data (GSE158142, only 8 dpf cells were used).

## Author contributions

B.H. and K.Q. designed research and supervised the overall direction of the research; Z.S. performed experiments with the assistance of L.Z., Y.S., H.Y., Y.C., L.S., A.H., and Z.Z.; Z.S. wrote the manuscript; B.H. and K.Q. revised the manuscript. All authors have read and agreed to the published version of the manuscript.

## Ethics approval and consent to participate

Not applicable

## Consent for publication

Not applicable

## References

1. Cadwell CR, Palasantza A, Jiang X, Berens P, Deng Q, Yilmaz M, Reimer J, Shen S, Bethge M, Tolias KF, et al: Electrophysiological, transcriptomic and morphologic profiling of single neurons using Patch-seq. Nat Biotechnol 2016, 34:199–203.

2. Cadwell CR, Scala F, Li S, Livrizzi G, Shen S, Sandberg R, Jiang X, Tolias AS: Multimodal profiling of single-cell morphology, electrophysiology, and gene expression using Patch-seq. Nat Protoc 2017, 12:2531–2553.

3. Bues J, Biocanin M, Pezoldt J, Dainese R, Chrisnandy A, Rezakhani S, Saelens W, Gardeux V, Gupta R, Sarkis R, et al: Deterministic scRNA-seq captures variation in intestinal crypt and organoid composition. Nat Methods 2022, 19:323–330.

4. Nieuwenhuis TO, Yang SY, Verma RX, Pillalamarri V, Arking DE, Rosenberg AZ, McCall MN, Halushka MK: Consistent RNA sequencing contamination in GTEx and other data sets. Nat Commun 2020, 11:1933.

5. Trapnell C: Defining cell types and states with single-cell genomics. Genome Res 2015, 25:1491–1498.

6. Kim HJ, Saikia JM, Monte KMA, Ha E, Romaus-Sanjurjo D, Sanchez JJ, Moore AX, Hernaiz-Llorens M, Chavez-Martinez CL, Agba CK, et al: Deep scRNA sequencing reveals a broadly applicable Regeneration Classifier and implicates antioxidant response in corticospinal axon regeneration. Neuron 2023, 111:3953–3969 e3955.

7. Cheng YH, Chen YC, Lin E, Brien R, Jung S, Chen YT, Lee W, Hao Z, Sahoo S, Min Kang H, et al: Hydro-Seq enables contamination-free high-throughput single-cell RNA-sequencing for circulating tumor cells. Nat Commun 2019, 10:2163.

8. Zhao P, Mondal S, Martin C, DuPlissis A, Chizari S, Ma KY, Maiya R, Messing RO, Jiang N, Ben-Yakar A: Femtosecond laser microdissection for isolation of regenerating C. elegans neurons for single-cell RNA sequencing. Nat Methods 2023, 20:590–599.

9. Marx V: Patch-seq takes neuroscience to a multimodal place. Nat Methods 2022, 19:1340–1344.

10. Arbabi K, Jiang Y, Howard D, Nigam A, Inoue W, Gonzalez-Burgos G, Felsky D, Tripathy SJ: Investigating microglia-neuron crosstalk by characterizing microglial contamination in human and mouse patch-seq datasets. iScience 2023, 26:107329.

11. Young MD, Behjati S: SoupX removes ambient RNA contamination from droplet-based single-cell RNA sequencing data. Gigascience 2020, 9.

12. Yang S, Corbett SE, Koga Y, Wang Z, Johnson WE, Yajima M, Campbell JD: Decontamination of ambient RNA in single-cell RNA-seq with DecontX. Genome Biol 2020, 21:57.

13. Korn H, Faber DS: The Mauthner cell half a century later: a neurobiological model for decision-making? Neuron 2005, 47:13–28.

14. Hu BB, Chen M, Huang RC, Huang YB, Xu Y, Yin W, Li L, Hu B: In vivo imaging of Mauthner axon regeneration, remyelination and synapses re-establishment after laser axotomy in zebrafish larvae. Exp Neurol 2018, 300:67–73.

15. Huang R, Chen M, Yang L, Wagle M, Guo S, Hu B: MicroRNA-133b Negatively Regulates Zebrafish Single Mauthner-Cell Axon Regeneration through Targeting tppp3 in Vivo. Front Mol Neurosci 2017, 10:375.

16. Hecker A, Anger P, Braaker PN, Schulze W, Schuster S: High-resolution mapping of injury-site dependent functional recovery in a single axon in zebrafish. Commun Biol 2020, 3:307.

17. Picelli S, Faridani OR, Bjorklund AK, Winberg G, Sagasser S, Sandberg R: Full-length RNA-seq from single cells using Smart-seq2. Nat Protoc 2014, 9:171–181.

18. van den Hurk M, Erwin JA, Yeo GW, Gage FH, Bardy C: Patch-Seq Protocol to Analyze the Electrophysiology, Morphology and Transcriptome of Whole Single Neurons Derived From Human Pluripotent Stem Cells. Front Mol Neurosci 2018, 11:261.

19. Lorent K, Liu KS, Fetcho JR, Granato M: The zebrafish space cadet gene controls axonal pathfinding of neurons that modulate fast turning movements. Development 2001, 128:2131–2142.

20. de Ceglia R, Ledonne A, Litvin DG, Lind BL, Carriero G, Latagliata EC, Bindocci E, Di Castro MA, Savtchouk I, Vitali I, et al: Specialized astrocytes mediate glutamatergic gliotransmission in the CNS. Nature 2023, 622:120–129.

21. Raj B, Farrell JA, Liu J, El Kholtei J, Carte AN, Navajas Acedo J, Du LY, McKenna A, Relic D, Leslie JM, Schier AF: Emergence of Neuronal Diversity during Vertebrate Brain Development. Neuron 2020, 108:1058–1074 e1056.

22. Mahar M, Cavalli V: Intrinsic mechanisms of neuronal axon regeneration. Nat Rev Neurosci 2018, 19:323–337.

23. Conforti L, Gilley J, Coleman MP: Wallerian degeneration: an emerging axon death pathway linking injury and disease. Nat Rev Neurosci 2014, 15:394–409.

24. Vargas ME, Barres BA: Why is Wallerian degeneration in the CNS so slow? Annu Rev Neurosci 2007, 30:153–179.

25. Feng Y, Yan T, Zheng J, Ge X, Mu Y, Zhang Y, Wu D, Du JL, Zhai Q: Overexpression of Wld(S) or Nmnat2 in mauthner cells by single-cell electroporation delays axon degeneration in live zebrafish. J Neurosci Res 2010, 88:3319–3327.

26. Halawani D, Wang Y, Ramakrishnan A, Estill M, He X, Shen L, Friedel RH, Zou H: Circadian clock regulator Bmal1 gates axon regeneration via Tet3 epigenetics in mouse sensory neurons. Nat Commun 2023, 14:5165.

27. Weng YL, An R, Cassin J, Joseph J, Mi R, Wang C, Zhong C, Jin SG, Pfeifer GP, Bellacosa A, et al: An Intrinsic Epigenetic Barrier for Functional Axon Regeneration. Neuron 2017, 94:337–346 e336.

28. Cho Y, Shin JE, Ewan EE, Oh YM, Pita-Thomas W, Cavalli V: Activating Injury-Responsive Genes with Hypoxia Enhances Axon Regeneration through Neuronal HIF-1alpha. Neuron 2015, 88:720–734.

29. Chen M, Long Q, Borrie MS, Sun H, Zhang C, Yang H, Shi D, Gartenberg MR, Deng W: Nucleoporin TPR promotes tRNA nuclear export and protein synthesis in lung cancer cells. PLoS Genet 2021, 17:e1009899.

30. Baggelaar MP, Maccarrone M, van der Stelt M: 2-Arachidonoylglycerol: A signaling lipid with manifold actions in the brain. Prog Lipid Res 2018, 71:1–17.

31. Xiang W, Shi R, Kang X, Zhang X, Chen P, Zhang L, Hou A, Wang R, Zhao Y, Zhao K, et al: Monoacylglycerol lipase regulates cannabinoid receptor 2-dependent macrophage activation and cancer progression. Nat Commun 2018, 9:2574.

32. Tanimura A, Yamazaki M, Hashimotodani Y, Uchigashima M, Kawata S, Abe M, Kita Y, Hashimoto K, Shimizu T, Watanabe M, et al: The endocannabinoid 2-arachidonoylglycerol produced by diacylglycerol lipase alpha mediates retrograde suppression of synaptic transmission. Neuron 2010, 65:320–327.

33. Chen M, Huang RC, Yang LQ, Ren DL, Hu B: In vivo imaging of evoked calcium responses indicates the intrinsic axonal regenerative capacity of zebrafish. FASEB J 2019, 33:7721–7733.

34. Vazquez-Sanchez S, Gonzalez-Lozano MA, Walfenzao A, Li KW, van Weering JRT: The endosomal protein sorting nexin 4 is a synaptic protein. Sci Rep 2020, 10:18239.

35. Jacobi A, Tran NM, Yan W, Benhar I, Tian F, Schaffer R, He Z, Sanes JR: Overlapping transcriptional programs promote survival and axonal regeneration of injured retinal ganglion cells. Neuron 2022, 110:2625–2645 e2627.

36. Bradke F: Mechanisms of Axon Growth and Regeneration: Moving between Development and Disease. J Neurosci 2022, 42:8393–8405.

37. Yook JS, Taxin ZH, Yuan B, Oikawa S, Auger C, Mutlu B, Puigserver P, Hui S, Kajimura S: The SLC25A47 locus controls gluconeogenesis and energy expenditure. Proc Natl Acad Sci U S A 2023, 120:e2216810120.

38. Jin D, Liu Y, Sun F, Wang X, Liu X, He Z: Restoration of skilled locomotion by sprouting corticospinal axons induced by co-deletion of PTEN and SOCS3. Nat Commun 2015, 6:8074.

39. Xie L, Yin Y, Benowitz L: Chemokine CCL5 promotes robust optic nerve regeneration and mediates many of the effects of CNTF gene therapy. Proc Natl Acad Sci U S A 2021, 118.

40. Chung D, Shum A, Caraveo G: GAP-43 and BASP1 in Axon Regeneration: Implications for the Treatment of Neurodegenerative Diseases. Front Cell Dev Biol 2020, 8:567537.

41. Gianola S, Rossi F: GAP-43 overexpression in adult mouse Purkinje cells overrides myelin-derived inhibition of neurite growth. Eur J Neurosci 2004, 19:819–830.

42. Aigner L, Arber S, Kapfhammer JP, Laux T, Schneider C, Botteri F, Brenner HR, Caroni P: Overexpression of the neural growth-associated protein GAP-43 induces nerve sprouting in the adult nervous system of transgenic mice. Cell 1995, 83:269–278.

43. Somasundaram P, Farley MM, Rudy MA, Stefanoff DG, Shah M, Goli P, Heo J, Wang S, Tran NM, Watkins TA: Coordinated stimulation of axon regenerative and neurodegenerative transcriptional programs by Atf4 following optic nerve injury. bioRxiv 2023.

44. Lindwall C, Kanje M: Retrograde axonal transport of JNK signaling molecules influence injury induced nuclear changes in p-c-Jun and ATF3 in adult rat sensory neurons. Mol Cell Neurosci 2005, 29:269–282.

45. Seijffers R, Mills CD, Woolf CJ: ATF3 increases the intrinsic growth state of DRG neurons to enhance peripheral nerve regeneration. J Neurosci 2007, 27:7911–7920.

46. Norsworthy MW, Bei F, Kawaguchi R, Wang Q, Tran NM, Li Y, Brommer B, Zhang Y, Wang C, Sanes JR, et al: Sox11 Expression Promotes Regeneration of Some Retinal Ganglion Cell Types but Kills Others. Neuron 2017, 94:1112–1120 e1114.

47. Song Y, Sretavan D, Salegio EA, Berg J, Huang X, Cheng T, Xiong X, Meltzer S, Han C, Nguyen TT, et al: Regulation of axon regeneration by the RNA repair and splicing pathway. Nat Neurosci 2015, 18:817–825.

48. Mason MRJ, van Erp S, Wolzak K, Behrens A, Raivich G, Verhaagen J: The Jun-dependent axon regeneration gene program: Jun promotes regeneration over plasticity. Hum Mol Genet 2022, 31:1242–1262.

49. Benarroch E: What Is the Role of Stathmin-2 in Axonal Biology and Degeneration? Neurology 2021, 97:330–333.

50. Moore DL, Blackmore MG, Hu Y, Kaestner KH, Bixby JL, Lemmon VP, Goldberg JL: KLF family members regulate intrinsic axon regeneration ability. Science 2009, 326:298–301.

51. English AW, Liu X, Mistretta OC, Ward PJ, Ye K: Asparagine Endopeptidase (delta Secretase), an Enzyme Implicated in Alzheimer’s Disease Pathology, Is an Inhibitor of Axon Regeneration in Peripheral Nerves. eNeuro 2021, 8.

52. Tedeschi A, Nguyen T, Puttagunta R, Gaub P, Di Giovanni S: A p53-CBP/p300 transcription module is required for GAP-43 expression, axon outgrowth, and regeneration. Cell Death Differ 2009, 16:543–554.

53. Du K, Zheng S, Zhang Q, Li S, Gao X, Wang J, Jiang L, Liu K: Pten Deletion Promotes Regrowth of Corticospinal Tract Axons 1 Year after Spinal Cord Injury. J Neurosci 2015, 35:9754–9763.

54. Duan X, Qiao M, Bei F, Kim IJ, He Z, Sanes JR: Subtype-specific regeneration of retinal ganglion cells following axotomy: effects of osteopontin and mTOR signaling. Neuron 2015, 85:1244–1256.

55. Bregman BS, Goldberger ME: Anatomical plasticity and sparing of function after spinal cord damage in neonatal cats. Science 1982, 217:553–555.

56. Zhou L, Kong G, Palmisano I, Cencioni MT, Danzi M, De Virgiliis F, Chadwick JS, Crawford G, Yu Z, De Winter F, et al: Reversible CD8 T cell-neuron cross-talk causes aging-dependent neuronal regenerative decline. Science 2022, 376:eabd5926.

57. Harel NY, Strittmatter SM: Can regenerating axons recapitulate developmental guidance during recovery from spinal cord injury? Nat Rev Neurosci 2006, 7:603–616.

58. Filbin MT: Recapitulate development to promote axonal regeneration: good or bad approach? Philos Trans R Soc Lond B Biol Sci 2006, 361:1565–1574.

59. Lu Y, Brommer B, Tian X, Krishnan A, Meer M, Wang C, Vera DL, Zeng Q, Yu D, Bonkowski MS, et al: Reprogramming to recover youthful epigenetic information and restore vision. Nature 2020, 588:124–129.

60. Hilton BJ, Bradke F: Can injured adult CNS axons regenerate by recapitulating development? Development 2017, 144:3417–3429.

61. Eaton RC, Farley RD: Development of the Mauthner Neurons in Embryos and Larvae of the Zebrafish, Brachydanio rerio. Copeia 1973, 1973:673–682.

62. Shao GJ, Wang XL, Wei ML, Ren DL, Hu B: DUSP2 deletion with CRISPR/Cas9 promotes Mauthner cell axonal regeneration at the early stage of zebrafish. Neural Regen Res 2023, 18:577–581.

63. Stern S, Hilton BJ, Burnside ER, Dupraz S, Handley EE, Gonyer JM, Brakebusch C, Bradke F: RhoA drives actin compaction to restrict axon regeneration and astrocyte reactivity after CNS injury. Neuron 2021, 109:3436–3455 e3439.

64. Shin JE, Geisler S, DiAntonio A: Dynamic regulation of SCG10 in regenerating axons after injury. Exp Neurol 2014, 252:1–11.

65. van den Brink SC, Sage F, Vertesy A, Spanjaard B, Peterson-Maduro J, Baron CS, Robin C, van Oudenaarden A: Single-cell sequencing reveals dissociation-induced gene expression in tissue subpopulations. Nat Methods 2017, 14:935–936.

66. O’Flanagan CH, Campbell KR, Zhang AW, Kabeer F, Lim JLP, Biele J, Eirew P, Lai D, McPherson A, Kong E, et al: Dissociation of solid tumor tissues with cold active protease for single-cell RNA-seq minimizes conserved collagenase-associated stress responses. Genome Biol 2019, 20:210.

67. Chen M, Xu Y, Huang R, Huang Y, Ge S, Hu B: N-Cadherin is Involved in Neuronal Activity-Dependent Regulation of Myelinating Capacity of Zebrafish Individual Oligodendrocytes In Vivo. Mol Neurobiol 2017, 54:6917–6930.

